# Specific F_1_ ATP synthase inhibition delivers transient mitochondrial stress for selective targeting of acute myeloid leukemia

**DOI:** 10.64898/2026.07.10.737821

**Authors:** Matthew T. Villaume, Haley E. Ramsey, Valeria Impedovo, Michael Davidson, Maria P. Arrate, Anand K. Singh, Yujin Lee, Anna Skwarska, Yara F. Almadani, Natalia Baran, Sovira Chaudhry, Benjamin J. Reisman, Eden G. TenBarge, Ming Jiang, Sarah Olmstead, Agnieszka E. Gorska, Zhiguo Zhao, Andrew J. Monteith, Peter M. Grace, Brian O. Bachmann, Marina Konopleva, Stefano Tiziani, Michael R. Savona

## Abstract

Targeting oxidative phosphorylation (OXPHOS) represents an attractive therapeutic strategy in acute myeloid leukemia, which exhibits exceptional dependence on mitochondrial respiration compared to normal hematopoietic cells. However, clinical attempts to exploit this vulnerability have been limited by on-target toxicity to healthy tissue. Here, we comprehensively compare the cellular consequences of inhibiting distinct nodes of the electron transport chain in AML. We demonstrate that selective inhibition of the F_1_ subunit of ATP synthase with EB2023 (ammocidin A) delivers an energetic stress to AML cells without the profound redox stress that characterizes complex I inhibition, preventing NAD⁺/NADH imbalance and allowing continued TCA cycling. Further, the duration of OXPHOS inhibition is transient in nature *in vivo*, a finding revealed through pharmacokinetic and serial pharmacodynamic monitoring of AMPK phosphorylation accompanied by OPA1-mediated mitochondrial structural remodeling that primes AML cells for BCL2 inhibitor synergy. EB2023 in combination with venetoclax demonstrates potent anti-AML activity across cell lines and patient-derived xenograft models at doses that spare normal hematopoietic progenitors and avoid the neuropathy and sustained detrimental systemic metabolic rewiring in healthy tissues associated with prior efforts to target OXPHOS. These findings establish F_1_-selective ATP synthase inhibition as a clinically actionable therapeutic strategy in AML and establish the duration of OXPHOS inhibition as a critical and previously underappreciated determinant of therapeutic index.

## Main

Metabolic reprogramming is a well-appreciated hallmark of cancer.^1^ This was first recognized by Otto Warburg in 1927 when he observed that tumors ferment glucose in the absence of oxygen,^2^ and has expanded with the observation that some cancer cells manipulate cellular energetics to rely on oxidative phosphorylation (OXPHOS - the electron transport coupled biosynthesis of ATP from ADP by ATP synthase) for bioenergetic and biosynthetic processes.^3–8^ Acute myeloid leukemia (AML) depends on OXPHOS and seems to have less metabolic flexibility than other cell types;^3,9–12^ and substantial evidence that the groundbreaking BCL2 inhibitor, venetoclax, imparts its AML lethality, at least in part, by suppressing AML OXPHOS, is crescendoing.^11,13^

Efforts to exploit OXPHOS inhibition therapeutically have been hampered by an inability to achieve selective toxicity in cancer cells without harming normal tissues. There are five main complexes within the electronic transport chain (ETC), each comprised of multiple subunit proteins and cofactors. While irreversible cessation of electron transport and OXPHOS is lethal to normal cells, partial or transitory inhibition has been studied, and recently, a novel inhibitor of complex I, IACS-010759 (I759), was revealed as a promising candidate in OXPHOS reliant cells and pursued in human study.^14^ Unfortunately, I759 led to significant on-target toxicity before an effective dose could be reached in clinical trials. This led to calls for a pause on further clinical development until a more comprehensive understanding OXPHOS inhibition as a treatment modality is known.^15,16^

The mitochondrial ATP synthase (complex V) is responsible for the terminal, ATP-generating step of OXPHOS and represents an alternative node for therapeutic intervention within OXPHOS. There are over a dozen protein subunits that comprise ATP synthase, and the tool F_O_-F_1_ inhibitor, oligomycin, canonically blocks the flow of protons through the membrane-imbedded F_O_ subunit, which in turn mechanically drives ATP synthesis by the catalytic F_1_-subunit. Unsurprisingly, in the 1980s, oligomycin was quickly found to lead to on-target cardiotoxicity in animal models, and was thus relegated to use as only a tool experimental compound in metabolic research.^17,18^ EB2023 (ammocidin A) is a unique glycomacrolide which we recently discovered to be a selective inhibitor of the F_1_ subunit of ATP synthase, and was safely administered to mice in AML xenograft models, with promising preliminary anti-AML activity.^19^ Here, using a systematic comprehensive approach, we demonstrate that F_1_ inhibition with EB2023 delivers a transient redox-balanced, energetic stress, disrupts mitochondrial structure in AML cells, and synergizes with low-dose venetoclax to sensitize venetoclax-resistant AML. These observations offer insight into the therapeutic index of selective ATP synthase inhibition and have implications for future drug development in this space.

## Results

As cancer cells activate divergent metabolic pathways from normal functioning cells, considerable effort has gone into understanding specific metabolic dependent liabilities in hopes of developing targeted metabolism-focused therapies.^20,21^ To examine metabolic dependencies across cancer cell lines, we queried the DepMap Public 25Q2 CRISPR-Cas9 essentiality dataset (Broad Institute, 2025; depmap.org/portal), which comprises genome-wide knockout screens across 1,186 cancer cell lines. Assessing four metabolism-related gene sets — glutamine metabolism, fatty acid oxidation, glycolysis, and OXPHOS — we found that cancer cell lines significantly vary in their dependence on metabolic pathways both between and within these categories. Many of the most essential metabolism-related genes originated from the OXPHOS gene set (Fig. 1a, see Methods). Within this OXPHOS gene set, we compared the relative dependencies between each component of the electron transport chain (complexes I-V) and again observed considerable heterogeneity. Specifically, though complex I remains the most frequently targeted ETC component in pre-clinical and clinical investigations,^22,23^ genetic depletion of complexes II, III and V show significantly greater mean anti-cancer effect (Fig. 1b, Extended Data Fig. 1a).

**Figure 1.**
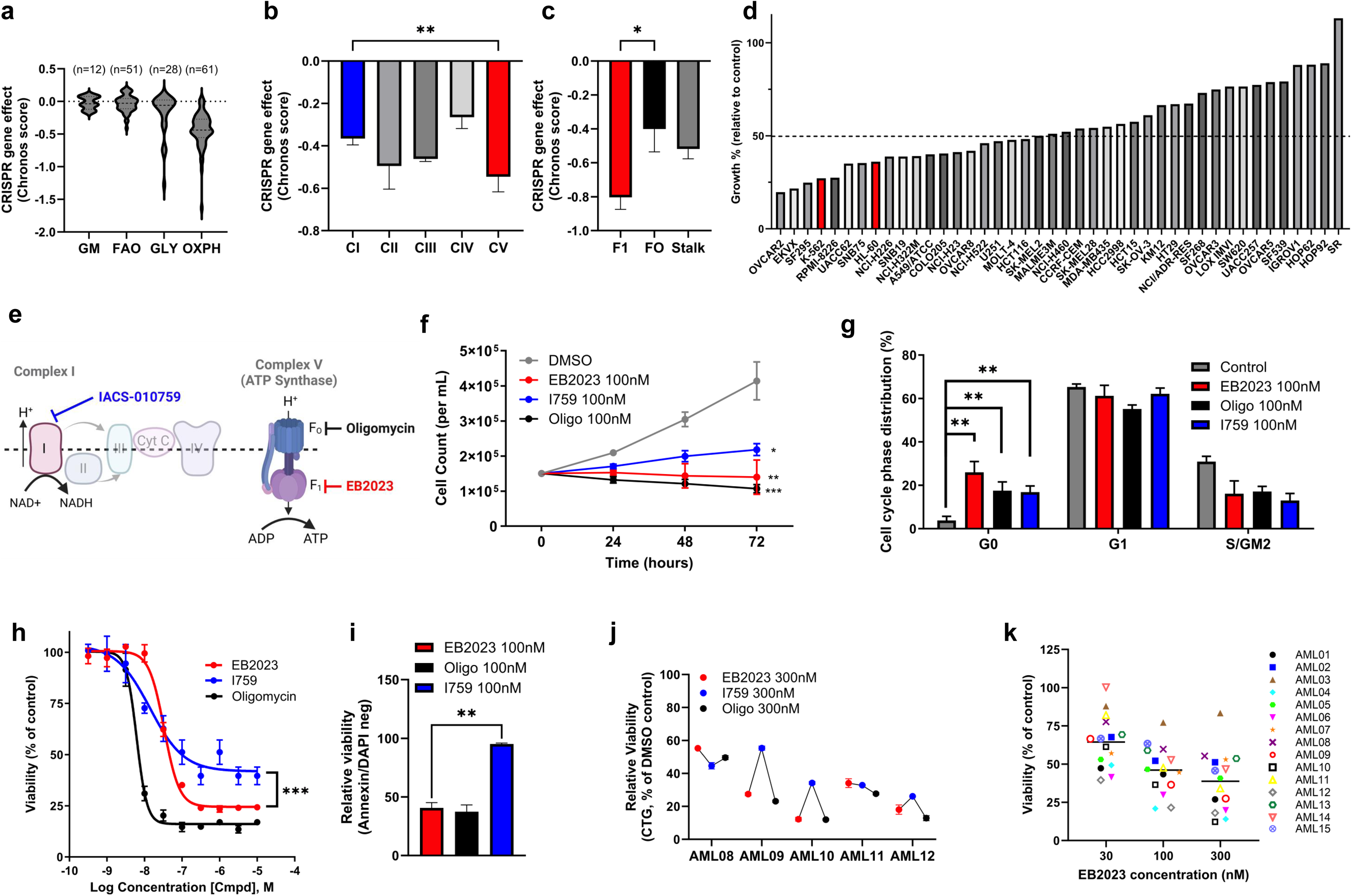
Selective genetic or chemical targeting of the F_1_ subunit of ATP synthase meaningfully impacts cancer cells. (a) Pan-cancer CRISPR-Cas9 gene dependency scores (Chronos) across four Gene Ontology metabolic gene sets — glutamine metabolism (GM), fatty acid oxidation (FAO), glycolysis (GLY), and oxidative phosphorylation (OXPH) — from the DepMap Public 25Q2 dataset (n = 1,186 cancer cell lines). Lower Chronos scores indicate greater essentiality. (b) Mean CRISPR-Cas9 dependency scores (Chrono) for nuclear-encoded ETC subunit genes grouped by complex (CI–CV) across 1,186 cancer cell lines from DepMap Public 25Q2. Mann-Whitney U test, Complex I vs Complex V. (c) Mean CRISPR-Cas9 gene dependency scores (Chronos) for ATP synthase subunit genes grouped by subcomplex (F_1_, F_O_, stalk) across 29 AML cell lines from DepMap Public 25Q2. Mann-Whitney U test, F_1_ vs F_O_. (d) Mean growth percent of NCI-60 cancer cell lines treated with apoptolidin A; AML cell lines highlighted in red. Dashed line indicates 50% growth. (e) Schematic of the mitochondrial electron transport chain illustrating the sites of action of I759, oligo, and EB2023. Created with BioRender.com. (f) Cell counts of MV-4-11 cells treated for 24, 48, and 72 h. (n = 3). Statistical comparisons were performed by two-way repeated measures ANOVA with Dunnett’s post-hoc test versus DMSO. (g) Cell cycle distribution of MV-4-11 cells treated for 24 hours (n = 3). G0 phase distributions were compared across treatment groups by one-way ANOVA with Dunnett’s post-hoc test versus control. (h) Dose-response curves in MV-4-11 cells assessed by CellTiter-Glo (CTG) viability assay at 48 hours (n = 6). Curves were fit by nonlinear regression using a four-parameter logistic model and Emax values were compared pairwise by Welch’s unpaired t-test. (i) Relative viability of MV-4-11 cells treated for 24 h, assessed by Annexin V/DAPI flow cytometry (n = 3). Early apoptotic fractions (Annexin V+/DAPI−) were compared between venetoclax and EB2023 + venetoclax by Welch’s unpaired t-test. (j) Relative viability (CTG) of primary AML patient samples treated for 72 hours (n = 3). Differences in viability across treatment conditions were assessed by Friedman test (χ²=4.80, p=0.091), with EB2023 and oligo showing greater viability reduction than I759 in 4 of 5 samples. (k) Viability (CTG) of 16 primary AML patient samples tested *ex vivo* across an EB2023 dose-response range. Data shown as mean ± SEM unless otherwise indicated. * p<0.05, ** p<0.01, *** p<0.001.

### Chemical inhibition of ATP synthase leads to anti-cancer lineage selectivity

Broad anti-cancer efficacy by targeting OXPHOS has been previously described but questions remain about how to target this pathway safely in humans.^24,25^ Many OXPHOS genes can be considered “pan-essential”, meaning in the public DepMap CRISPR-Cas9 data set, these genes rank among the most depleting knockouts in >90% of cancer cell lines screened, with no lineage specificity, and therefore imply they are not likely be a safe drug target in humans.^26^ Each ETC complex is encoded by multiple pan-essential genes (i.e. NDUFS5, ATP5MF, and ATP5F1B; Extended Data Fig. 1b). While genetic depletion of complex V (ATP synthase) had broad, indiscriminate anti-cancer effects, pharmacological/chemical inhibition of ATP synthase had a more selective effect on hematopoietic cancer cell lines. To this end, the canonical F_O_-F_1_ inhibitor oligomycin had a broad range of anti-cancer efficacy but was most potent against myeloid malignancy cell lines (Extended Data Fig. 1c). This anti-cancer lineage enrichment was comparable to the selectivity seen with standard-of-care venetoclax therapy against myeloid cell lines (Extended Data Fig. 1d). This illustrates that chemically targeting ATP synthase with a suitable therapeutic window may be feasible despite CRISPR-Cas9 screening results that suggest otherwise. In contrast to this, the selective complex I inhibitor, I759, did not show any lineage enrichment for any cancer type (Extended Data Fig. 1e).

### F_1_-subunit of ATP synthase can be chemically targeted with EB2023

Further sub-classification of ATP synthase gene dependency in cancer and specifically AML cell lines revealed that the knockout of genes encoding the F_1_-subunit of ATP synthase had a greater effect on cancer cell line growth than did those encoding the F_O_-subunit, particularly in AML cell lines (Fig. 1c). Consistent with this finding, chemical inhibition of this target with single dose testing of a F_1_-targeting glycomacrolide natural product apoptolidin against a NCI60 cancer cell line panel revealed variable activity across tumors, with considerable activity in AML cell lines (Fig. 1d). EB2023 is the most potent member of this F1-subunit targeting family of glycomacrolide natural products,^27^ so we further investigated the most promising lineages identified in the NCI60 panel by testing EB2023 against a larger panel of AML, multiple myeloma, melanoma and breast cancer cell lines and observed 50-100% reductions in viability after 72 hours of treatment in most lines with IC_50_ values ranging from 6-41nM (CellTiter-Glo® (CTG), Extended Data Fig. 2a-d). We compared viabilities as measured by CTG, where relative luminescence approximates ATP quantities, with an orthogonal assay which measures cell protease activity (CellTiter-Fluor®) and demonstrated concordant results (Extended Data Fig. 2e-g).

The differences observed in cancer gene dependency within the ETC complex and even individual subunits of these complexes motivated us to pursue a direct comparison of chemical inhibitors of these targets. We tested a complex I inhibitor (I759) and two mechanistically distinct ATP synthase inhibitors (oligomycin and EB2023) across an array of cell growth and viability assays to elucidate whether anti-AML activity varied by ETC target (Fig. 1e). Cell growth was inhibited by all three inhibitors (Fig. 1f). While all three drugs decreased cell cycling, as evidenced by increased G0 and decreased G2/S/M phase cells (Fig.1g), the ATP synthase inhibitors more potently induced cell death in AML cell lines as measured by CTG (Fig. 1h, Supplemental Fig. 1) and Annexin V/DAPI staining (Fig. 1i). We assessed cell viability following treatment with EB2023, I759, or oligomycin at 300 nM across five patient-derived AML specimens. All three inhibitors reduced viability relative to DMSO control and Friedman test across the three treatment conditions did not reach statistical significance (χ²=4.80, p=0.091, Fig. 1j, Supplemental Fig. 1a-f).

### EB2023 is a selective F_1_ ATP synthase inhibitor with broad anti-AML activity which disrupts mitochondrial respiration

We sought to expand efficacy assessments of EB2023 in AML patient samples *ex vivo* (Fig 1k) and 10 AML cell lines (Extended Data Fig. 3a-b) with a range of EB2023 concentrations for 48 hours. F_1_ ATP synthase inhibition reduced the cell growth kinetics in a dose-dependent fashion in all samples tested with IC_50_ less than 50nM in 9 out of 10 cell lines and <100nM in a majority of patient samples, with viability reductions occurring well below safely achievable doses *in vivo*.^19^ We observed dose-dependent reductions in basal oxygen consumption rate in MV-4-11 cells and significant reductions were seen in 3 of our 4 cell lines at a dose of 100nM (Extended Data Fig. 3c-d).

Suppression of OXPHOS and the TCA cycle is known to relieve negative feedback mechanisms on glycolysis, which is a compensatory metabolic escape seen with many inhibitors of OXPHOS.^14,28,29^ Accordingly, enhanced EB2023 cytotoxicity occurred when AML cells were cultured in galactose media when compared to human plasma like media (HPLM; Extended Data Fig. 3e). Pyruvate dehydrogenase kinase 1 (PDK1) has been described as a “glycolytic gatekeeper” in AML with lower expression levels defining OXPHOS-driven leukemias with decreased glycolytic reserve.^30^ Using PDK1 expression data from DepMap Public 25Q2 data set, AML cell lines most sensitive to EB2023 were found to have significantly lower expression levels of PDK1 (Extended Data Fig. 3f). This finding is again consistent with the proposed mechanism of EB2023’s anti-AML effect being driven by suppression of mitochondrial respiration.

### Nanomolar dosing of ATP synthase inhibitors causes a transient uncoupling of AML respiration prior to halting respiration

Towards better understanding these discrepant anti-AML effects, we next assessed how this panel of OXPHOS inhibitors affect AML metabolism. We selected 100nM as the initial comparison concentration for all three inhibitors as this was the lowest dose needed to achieve maximal anti-AML effect for all three drugs across all samples tested (Supplemental Fig. 1). All three inhibitors suppressed oxygen consumption as measured by Seahorse XF analysis (OCR) after 24 hours of exposure in MV-4-11 (Fig. 2a-b) and Molm-13 cells (Extended Data Fig. 4a). Complex I inhibition had a greater effect on spare respiratory capacity in both cell lines consistent with its mechanism of action disrupting NADH-driven proton shuttling into the intermembrane space. While I759 induced an anticipated increase in glycolysis (ECAR) in MV-4-11 cells, neither of the ATP synthase inhibitors lead to increases in glycolysis after 24 h of drug exposure in this cell line (Fig. 2c), while in the less sensitive Molm-13 line, all three drugs triggered increases in ECAR (Extended Data Fig. 4b). To further investigate this result, we measured OCR and ECAR immediately after addition of 100nM of each drug and saw that oligomycin and EB2023 caused a transient increase in OCR and did not affect ECAR over the first 4 hours (Fig. 2d). This finding was unanticipated as oligomycin is a well-characterized tool compound for disrupting OXPHOS by inhibiting ATP synthase at micromolar concentrations; however, there are reports that this effect may be dose dependent and oligomycin-induced proton uncoupling has been observed in certain experimental contexts.^31^ To this end, both ATP synthase inhibitors caused an immediate cessation in oxygen consumption and induction of glycolysis when used at micromolar concentrations in the same experiment (Extended Data Fig. 4c), highlighting that low-dose ATP synthase inhibition has unique effects on AML respiration that are suggestive of an uncoupling of ATP production from respiration at early timepoints. Consistent with uncoupled mitochondria activity, the levels of mitochondrial reactive oxygen species showed decreases rather than the increases seen with I759 (Extended Data Fig. 4d), suggesting that ATP synthase inhibitors at nanomolar doses allow the continued flow of electrons from NADH across Complexes I-IV. Notably, there was no concomitant decrease in mitochondrial membrane potential at these early time points with either ATP synthase inhibitor at nano-molar concentrations, and reductions in membrane potential, as has been previously described to occur rapidly with oligomycin treatment, occurred only with long term exposure of EB2023 (Extended Data Fig. 4e-f).^32^ In summary, at 100 nM concentrations, which are 5 - 10 fold their IC_50_ concentration, both EB2023 and oligomycin produced a transient increase in OCR without a corresponding increase in ECAR, indicating that Fo-mediated proton conductance remains intact under conditions of selective F₁ and Fo inhibition, and that compensatory glycolysis induction is avoided in certain cases.

**Figure 2.**
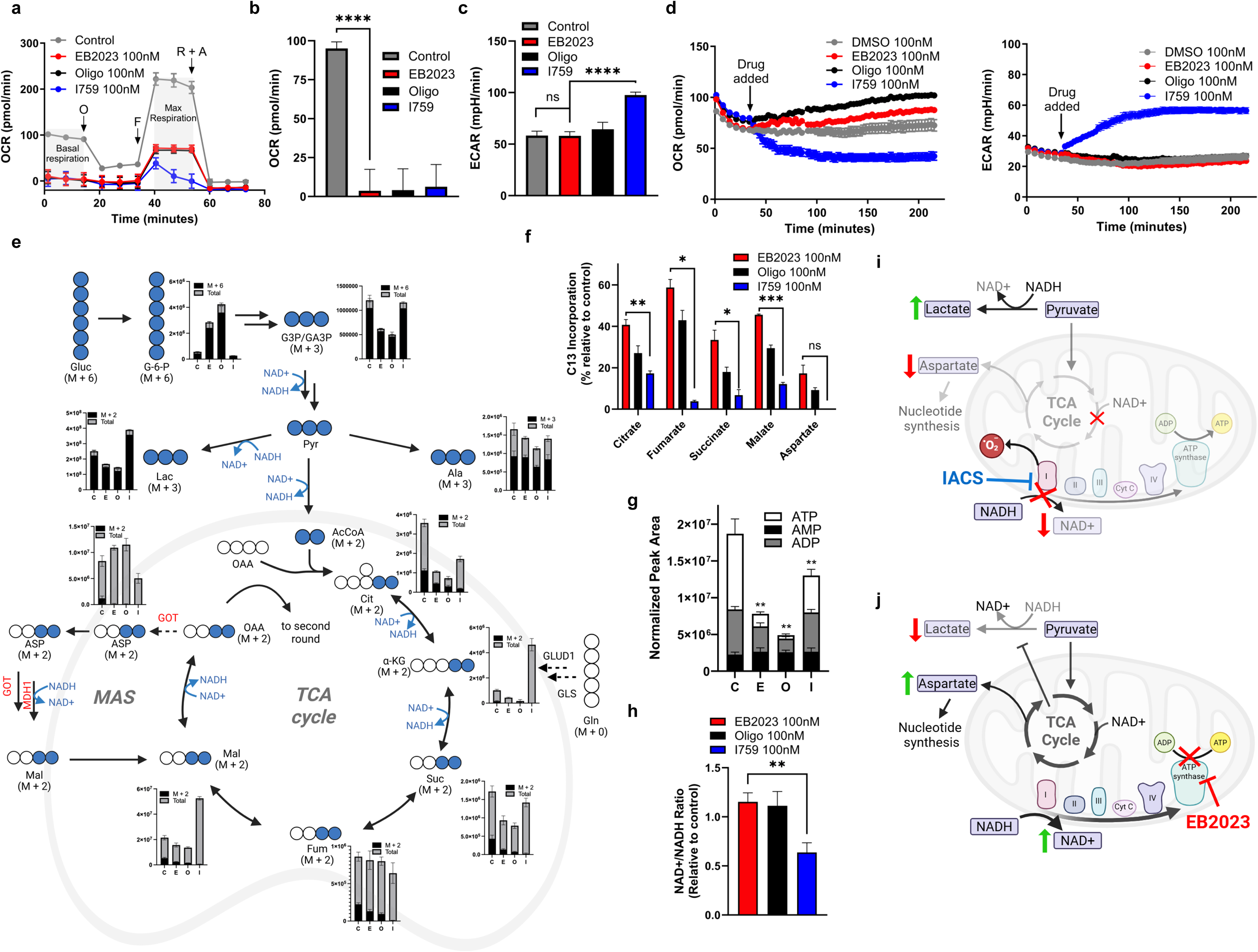
Nanomolar ATP synthase inhibition uncouples AML respiration and imposes energetic stress while preserving NAD⁺ redox state. (a) OCR measured during Seahorse XF Mito Stress Test in MV-4-11 cells pretreated for 24 hours; representative of n = 3 independent experiments. O, oligomycin; F, FCCP; R+A, rotenone and antimycin A. (b) Basal respiration of MV-4-11 cells after 24 hours of treatment quantified from the Seahorse XF Mito Stress Test. Basal respiration was compared between DMSO and each inhibitor by Welch’s unpaired t-test. (c) Extracellular acidification rate of MV-4-11 cells after 24 hours of treatment with DMSO, EB2023 (100 nM), oligo (100 nM), or I759 (100 nM), measured by Seahorse XF Glycolysis Stress Test (Agilent). Comparisons performed by Welch’s unpaired t-test. (d) Real-time OCR (left) and ECAR (right) of MV-4-11 cells following injection of indicated drug measured by Seahorse XF analyzer. (e) Stable isotope tracing schematic of [U-¹³C₆]-glucose metabolism in MV-4-11 cells treated with DMSO (C), EB2023 (E), oligo (O), or I759 (I) for 24 hours. Bar graphs depict labeled (M+x, black) and total metabolite abundances (grey) measured by LC-MS. Arrows indicate key metabolic reactions; NAD⁺/NADH cofactor utilization is annotated at relevant steps. (f) ¹³C-labeled fraction of TCA cycle intermediates as a percentage of total metabolite abundance in MV-4-11 cells treated as in (e) (n = 4). ¹³C isotope incorporation was compared across treatment groups by two-way ANOVA with treatment and metabolite as factors; both the main effect of treatment and the treatment × metabolite interaction were significant (both p<0.0001). Post-hoc comparisons to I759 were performed using Welch’s unpaired t-test with Holm-Bonferroni correction for multiple comparisons. (g) Total abundance of ATP, ADP, and AMP in MV-4-11 cells treated with DMSO (C), EB2023 (E), oligo (O), or I759 (I) for 24 hours, measured by LC-MS (n = 4). ATP normalized peak areas were compared by Welch’s one-way ANOVA (F=62.2, p<0.0001) with Holm-Bonferroni corrected post-hoc Welch’s t-tests versus control. (h) NAD⁺/NADH ratio in MV-4-11 cells treated for 4 hours, measured by colorimetric assay (Abcam, n = 4). Values normalized to DMSO control. Comparisons to DMSO performed by Welch’s unpaired t-test. (i–j) Schematics summarizing the metabolic consequences of Complex I inhibition by I759 (i) versus F_1_ ATP synthase inhibition by EB2023 (j). Created with BioRender.com. Error bars represent mean ± SEM throughout. ns = not significant, *p<0.05, **p<0.01, ***p<0.001, ****p<0.0001.

### EB2023 delivers an energetic stress without a redox stress from imbalance of NAD^+^/NADH to AML cells while allowing persistent TCA cycling of glucose carbon

We next compared the inhibitors across a targeted metabolomics panel and stable-isotope tracing experiments. Two AML cell lines with divergent basal respiration rates (MV-4-11, Fig. 2e; and Molm-13, Extended Data Fig. 4g) were exposed to each OXPHOS inhibitor for 24 h and the resultant cell lysates were subjected to liquid chromatography-mass spectrometry (LC-MS) analysis. The total abundance of TCA-related metabolites and the relative abundance of [¹³C]-glucose-derived isotopically labeled metabolites showed consistent differences between the complex I and the ATP synthase inhibitors. All inhibitors resulted in decreased incorporation of labeled carbon into TCA cycle intermediates (Fig. 2f). However, there was a higher incorporation of glucose-derived carbons in the TCA cycle upon treatment with ATP synthase inhibitors compared to complex 1 inhibitor in both cell lines. All three inhibitors significantly reduced total ATP and ATP/ADP ratio by LCMS analysis in MV-4-11 cells (Fig. 2g), though Molm-13 cells were able to better maintain ATP levels in presence of the ATP synthase inhibitors (Extended Data Fig. 4h). Continued TCA-cycling with ATP synthase inhibitors resulted in preserved synthesis of the nucleotide precursor aspartate (Fig. 2f), and supplementation of exogenous aspartic acid partially rescued MV-4-11 cell growth in the presence of I759 but had no effect in the presence of EB2023 (Extended Data Fig. 4i). This is consistent with prior observations implicating nucleotide precursor depletion in I759-mediated growth suppression,^14^ which does not occur with treatment of EB2023. Additionally, there were significant increases in the total levels of some unlabeled TCA cycle intermediates alpha-ketoglutarate and malate seen after inhibition of complex I, that was not observed with ATP synthase inhibition; this phenomenon has previously been attributed to compensatory upregulation of glutamine metabolism in response to complex I inhibition (Fig. 2e, Extended Data Fig. 4g).^14^

We hypothesized that a relative preservation of NAD^+^ in the presence of ATP synthase inhibitors, primarily generated from uninhibited complex I NADH oxidation, could explain both the uninterrupted oxygen consumption and preserved functioning of TCA cycle reactions. Consistent with this, MV-4-11 cells showed unchanged NAD^+^/NADH ratios after 4 hours of treatment with ATP synthase inhibitors, while this ratio was rapidly reduced with complex I inhibition (Fig. 2h). This preserved oxidation state, along with more moderate reductions in TCA metabolites, does not trigger rapid lactic acid production in the cytosol as was seen with complex I inhibition (Fig. 2e). In summary, while all three OXPHOS inhibitors deliver an energetic stress to AML cells, ATP synthase inhibitors do this without causing a redox stress with shared metabolic phenotypes of the three drugs being related to energy stress adaptations while differences can be explained by preserved NAD^+^-dependent TCA enzyme reactions (Fig. 2i-j).

### F_1_ ATP synthase inhibition triggers distinct cell stress response pathways and disrupts mitochondrial structure via perturbations of OPA1

To characterize transcriptional responses to the OXPHOS inhibitors, we performed RNA sequencing and gene set enrichment analysis (GSEA) after dosing MV-4-11 (more sensitive) and THP-1 (less sensitive) cells with EB2023, I759, or oligomycin for 24 hours (Fig. 3a-d, Extended Data Fig 5a). All three compounds triggered increased expression of integrated stress response (ISR) target genes (ATF4, DDIT4, TRIB3; Fig. 3a-c), with EB2023 and oligomycin both triggering the heme deficiency (HRI) pathway, previously identified to be operative in response to ATP synthase inhibition (Fig. 3d). All three compounds suppressed various cell cycle pathways, including G1/S transition, S phase, and mitotic progression (Fig. 3d, Extended Data Fig. 5a) in both cell lines regardless of sensitivity. The changes observed in cell cycling and numerous changes in critical modulators of metabolism (PHGDH, SQLE, FASN, SCD), are consistent with ATP depletion triggering increases in AMPK activity. Dose-dependent phosphorylation of AMPK (Extended Data Fig. 5b) and eIF2α of the integrated stress response along with their downstream effectors (ACC, ATF4 respectively) were observed at this timepoint (Extended Data Fig. 5c). These findings are largely consistent with prior data of leukemia cells treated with I759, in which AMPK signaling and the integrated stress response have been previously implicated and likely represent shared characteristics of OXPHOS inhibition.^14,28,33^ Interestingly, the less EB2023-sensitive cell line, THP-1, upregulated genes related to import of amino acids in response to all three drugs, suggesting a mechanism of metabolic adaptation to OXPHOS inhibition that MV-4-11 cannot replicate (Extended Data Fig. 5a).

**Figure 3.**
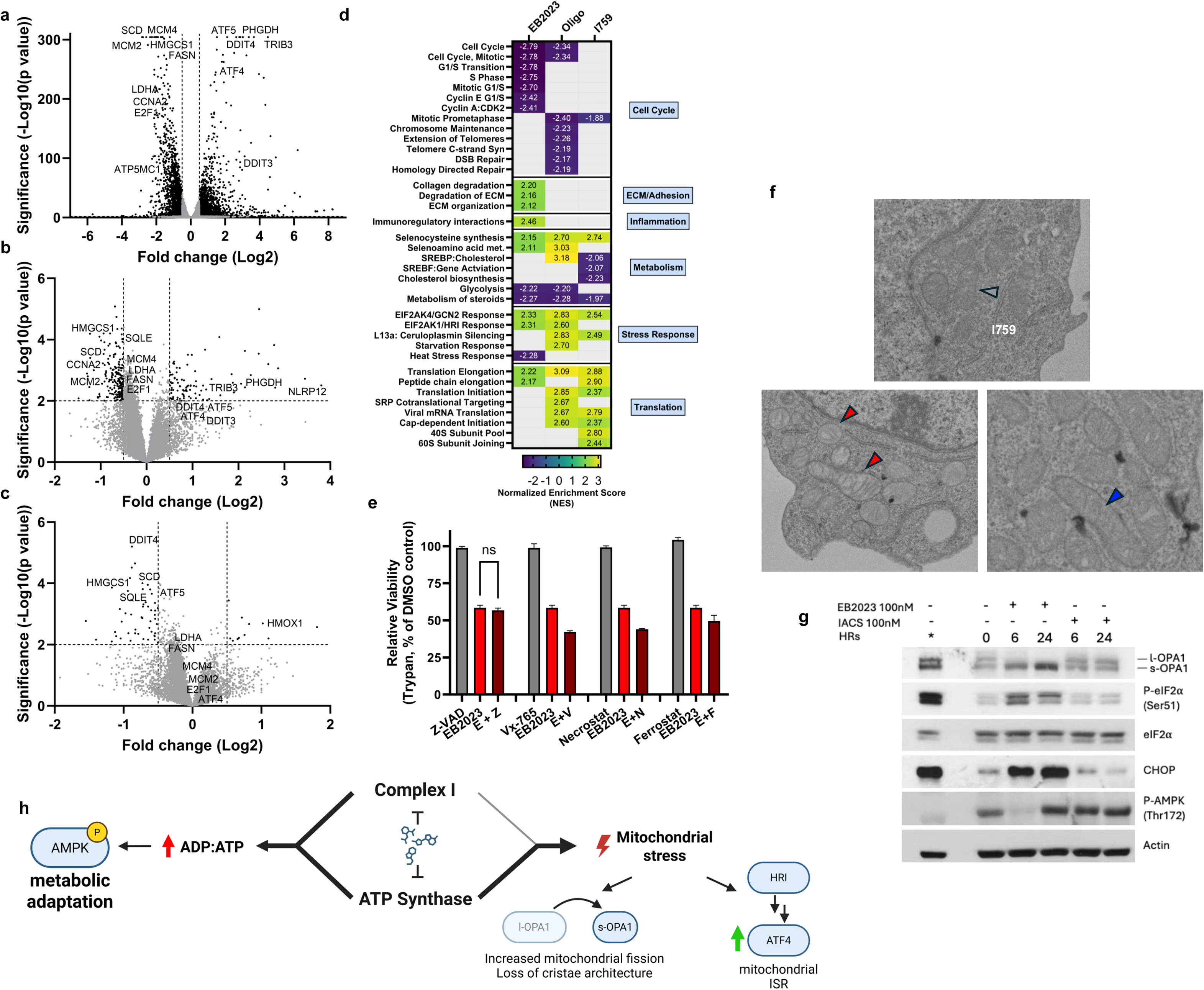
OXPHOS inhibition triggers AMPK activation but ATP synthase inhibition delivers a unique stress to mitochondria and leads to caspase-independent cell death. (a–c) Volcano plots showing differentially expressed genes in MV-4-11 cells treated with EB2023 (a), oligomycin (b), or I759 (c) at 100 nM for 24 hours versus DMSO control. The x-axis represents log₂ fold change and the y-axis represents −log₁₀(adjusted p-value). Dashed vertical lines indicate log₂ fold change thresholds; dashed horizontal line indicates the significance threshold. Selected genes involved in metabolic and stress response pathways are labeled. (d) Heatmap of normalized enrichment scores (NES) from gene set enrichment analysis (GSEA) of Reactome gene sets in MV-4-11 cells treated with EB2023, oligomycin, or I759 at 100 nM for 24 hours compared to DMSO control. Gene sets are grouped by biological category. Color scale represents NES; only gene sets meeting significance thresholds for at least one comparison are shown. (e) Relative viability (trypan blue exclusion) of MV-4-11 cells treated with EB2023 (100 nM) alone or in combination with inhibitors of caspase-dependent apoptosis (Z-VAD-FMK, 10 µM), caspase-1 (Vx-765, 1 µM), necroptosis (Necrostatin-1, 1 µM), or ferroptosis (Ferrostatin-1, 5 µM) for 48 hours. Comparisons to EB2023 alone performed by Welch’s unpaired t-test. (f) Representative TEM images of MV-4-11 mitochondria in DMSO control, EB2023-treated, and I759-treated cells at 24 hours. Grey arrowheads indicate untreated mitochondrial morphology; red arrowheads indicate round mitochondria with loss of cristae architecture; blue arrowheads indicate elongated mitochondria with tight cristae. (g) Immunoblot analysis of OPA1 (long/short isoforms), phospho-eIF2α (Ser51), total eIF2α, CHOP, phospho-AMPK (Thr172), and actin (loading control) in MV-4-11. (h) Schematic summarizing the divergent downstream consequences of Complex I inhibition versus ATP synthase inhibition in AML cells, highlighting shared AMPK activation and divergent effects on mitochondrial stress signaling, OPA1 proteolysis, cristae architecture, and ISR activation. Created with BioRender.com. Error bars represent mean ± SEM throughout. ns = not significant, *p<0.05, **p<0.01, ***p<0.001, ****p<0.0001.

To better understand the mode of cell death and ultrastructural changes in response to OXPHOS inhibition we pursued transmission electron microscopy (TEM) of MV-4-11 cells treated with OXPHOS inhibitors. TEM revealed that EB2023 induced significant vacuolization of the cytoplasm and whole-cell morphology changes consistent with loss of membrane integrity and organelle swelling without apoptotic nuclear changes (Extended Data Fig. 5d). Similar results were observed with oligomycin treatment though to a lesser extent (Extended Data Fig. 5e). In contrast, I759 treated cells had far fewer morphologic changes at 24h with significant changes only occurring at 72h of treatment (Extended Data Fig. 5f-g). EB2023 led to loss of cell membrane integrity in both MV-4-11 and Molm-13 (Extended Data Fig. 5h-i) as measured by trypan exclusion but did not induce caspase activity consistently in these different cell lines (Extended Data Fig. 5j). The pan-caspase inhibitor Z-VAD-FMK was unable to reduce EB2023-induced cell death (Fig. 3e), further implicating a caspase-independent cell death mechanism. Inhibitors targeting key components of other programmed necrotic cell death pathways, such as necroptosis (RIPK1 inhibitor, Necrostatin-1), pyroptosis (caspase-1 inhibitor, Vx-765), and ferroptosis (radical-trapping antioxidant, Ferrostatin-1) also failed to prevent cell death.

Examining mitochondrial structure from TEM images revealed that AML mitochondria from cells treated with EB2023 were more electron-lucent (Extended Data Fig. 5k) and rounder (Extended Data Fig. 5l) with more significant distortions in cristae structure (Fig. 3f). Cristae deformations and increased mitochondrial roundness are seen with severe mitochondrial stress and specifically have been correlated with changes in OPA1 protein proteolytic cleavage from its long form (l-OPA1) to its short form (s-OPA1).^34^ While both EB2023 and I759 triggered the phosphorylation of AMPK by 24 hours as anticipated, only ATP synthase inhibition caused evidence of downstream ISR activation and an increase in s-OPA1 to l-OPA1 ratio (Fig. 3g). Redistribution to s-OPA1 has been shown to result in the fragmentation of the mitochondrial network (loss of cristae structure seen with EB2023), loss of mitochondrial fusion capacity (evidenced by greater roundness in EB2023 compared to I759), and reduced ATP production, which can induce cellular stress and apoptosis.^35,36^ Complex I inhibition with I759 produces a relatively isolated energy-stress signature (AMPK activation), whereas EB2023 produces a broader mitochondrial stress program involving OPA1 processing and ISR activation. This suggests ATP synthase inhibition is not merely a weaker version of complex I inhibition, but a qualitatively distinct mitochondrial insult (Fig. 3h). These same changes in mitochondrial structure and OPA1 have also been correlated with BCL2 dependence, prompting further investigation into EB2023’s combination with the BCL2 inhibitor venetoclax (VEN).^34^

### EB2023 synergistically induces apoptosis with venetoclax even at sublethal monotherapy doses

Molm-13 is a FLT3-mutated cell line known to be OXPHOS-driven^30^ but was the least sensitive cell line screened against I759.^14^ EB2023 slowed but does not stop Molm-13 cell growth *in vitro* (Fig. 4a) and incompletely inhibited OCR, with >50% of baseline OCR occurring in the presence of 100nM EB2023 (Fig. 4b). Despite this, EB2023 demonstrated potent synergy with venetoclax against Molm-13 and most other AML cell lines (ZIP Synergy score^37,38^, Fig. 4c, Extended Data Fig. 6a). Neither 100nM venetoclax nor 100nM EB2023, as single agents, induce significant apoptosis in either Molm-13 or THP-1 cell lines, but their combination increased caspase 3/7 activity (Extended Data Fig. 6b) and annexin V staining (Fig. 4d). Unlike with EB2023 monotherapy, combination-induced cell death was partially protected by the addition of the pan-caspase inhibitor Z-VAD-FMK (Extended Data Fig. 6c), further supporting the notion of caspase-dependent apoptosis with the combination. Similar increases in apoptotic cell death were observed with the combination against AML blasts in a panel of AML patient samples (Extended Data Fig. 6d).

**Figure 4.**
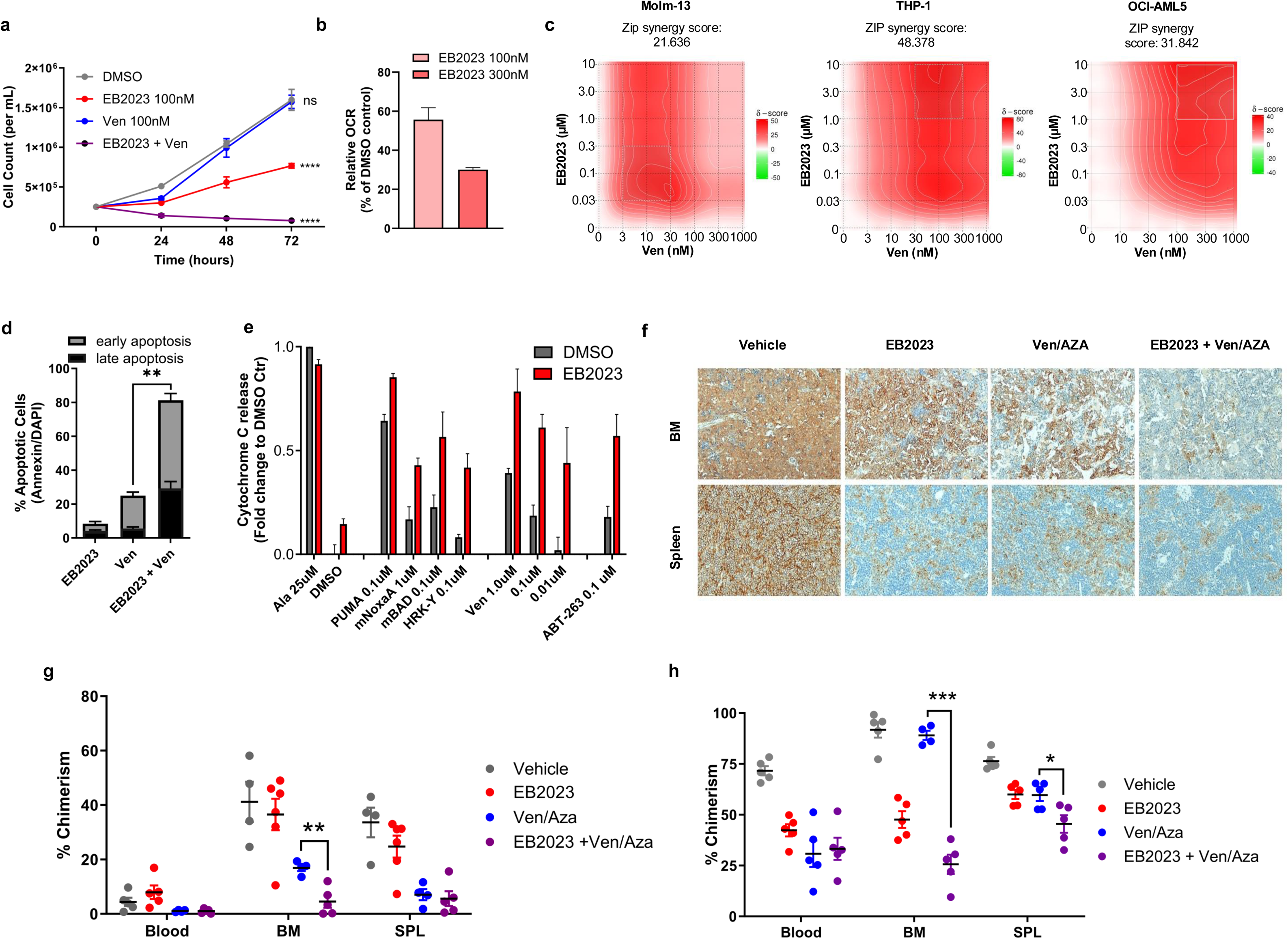
EB2023 primes AML mitochondria for BCL2 dependence and overcomes venetoclax resistance at sub-lethal doses. **(a)** Cell counts of Molm-13 cells treated for 72 hours with EB2023, venetoclax (Ven) or combination of both, assessed by automated cell counting (Countess 3) (n = 3). Statistical comparisons were performed by two-way repeated measures ANOVA with Dunnett’s post-hoc test versus DMSO. **(b)** Basal OCR of Molm-13 cells treated for 24 hours, measured by Seahorse XF Mito Stress Test and expressed as percent of DMSO control (n = 6). Comparisons to DMSO performed by Welch’s unpaired t-test (p <0.01). **(c)** ZIP synergy score matrices for EB2023 (10 nM–10 µM) and venetoclax (Ven; 1 nM–1 µM) across three AML cell lines (Molm-13, THP-1, OCI-AML5), assessed by CellTiter-Glo viability assay at 48 hours. ZIP synergy scores are indicated for each cell line. Warm colors indicate synergistic interactions; cool colors indicate antagonism. ZIP synergy score >10 indicates synergy.^37^ **(d)** Percentage of early (Annexin V+/DAPI−) and late (Annexin V+/DAPI+) apoptotic Molm-13 cells treated with EB2023 (100 nM), venetoclax (30 nM), or their combination for 18 hours (n = 3). Early apoptotic fractions were compared between venetoclax and EB2023 + venetoclax by Welch’s unpaired t-test. **(e)** BH3 profiling of THP-1 cells pretreated with DMSO or EB2023 (100 nM) for 24 hours (n = 3). **(f)** Representative hCD45 immunohistochemistry of bone marrow (BM) and spleen sections from MV-4-11 CDX NSGS mice treated with vehicle, EB2023, venetoclax + azacitidine (Ven/AZA), or EB2023 + Ven/AZA (n = 6 per group). **(g-h)** Human AML chimerism (huCD45/huCD45 +moCD45 cells by flow cytometry) in blood, BM, and spleen of Molm-13 CDX (g) and AML06 PDX (h) NSGS mice (n = 6 per group) treated with vehicle, EB2023, Ven/AZA, or EB2023 + Ven/AZA. Comparisons performed by Welch’s unpaired t-test. All bar and dot plots represent mean ± SEM *p<0.05, **p<0.01, ***p<0.001, ****p<0.0001.

To better understand this synergy, we performed BH3 profiling analysis on THP-1 cells which are *TP53* and *NRAS* mutated and relatively venetoclax insensitive (IC_50_ 1.85µM).^39,40^ EB2023 pretreatment potently primed THP-1 cells to BCL2 and BCLxL dependence, as evidenced by significant cytochrome c release in response to BCL2/BCLxL-specific BAD and BCLxL-specific HRK peptides, as well as to the BCL2 inhibitor venetoclax and dual BCL2/BCLxL inhibitor, ABT-263 (Fig. 4e, Extended Data Fig. 6e). To a lesser extent, EB2023 sensitized cells to MCL1-specific peptides such as NOXA and MS1 or MCL1 inhibitor S63845. Consistent with this BCL2/BCLxL priming mechanism, MCL1 protein abundance was shown to decrease with EB2023 exposure (Extended Data Fig. 6f), though this could be due to initiation of cell death processes rather than an ISR driven epigenetic change as has been proposed with azacitidine.^41,42^ Cells exposed to EB2023 and venetoclax combination showed rapid loss of mitochondrial membrane potential (Extended Data Fig. 6g) and complete abrogation of the persistent oxygen consumption seen with either monotherapy (Extended Data Fig. 6h).

### EB2023 synergizes with low-dose venetoclax in venetoclax-resistant models *in vivo*

Based on the synergistic effect of EB2023 and venetoclax combination *in vitro*, we tested the addition of EB2023 to the standard of care AML regimen of venetoclax and azacitidine *in vivo*. Multiple xenograft transplantation models of AML were assessed in NSGS mice, including 2 cell line-derived (CDX, MV-4-11 and Molm-13) and 2 patient-derived xenograft models (PDX). After establishing disseminated leukemia, mice were treated with either: EB2023 (0.1 mg/kg i.p., for 5 days on/2 days off continuously), low-dose venetoclax (20mg/kg oral gavage, for 5 days on/2 days off continuously), azacitidine (1.5 mg/kg i.p. daily for 3 days), or the triplet combination. For the CDX models, mice were treated for 21 days then sacrificed and human CD45+/CD33+ cells were measured by flow cytometry to assess chimerism. MV-4-11 transplanted mice showed modest reductions in AML burden with both EB2023 and ven/aza at these doses, but the triplet combination showed significant reductions in all compartments by flow cytometry and IHC CD45 immunohistochemistry (Fig. 4f and Extended Data Fig. 6i-j). In the Molm-13 CDX model, EB2023 showed minimal single agent activity but significantly enhanced clearance of AML cells by the triplet regimen from the bone marrow, as predicted by *in vitro* experiments (Fig. 4g).

For the PDX models, we chose a *NPM1*/*FLT3-ITD*-mutated AML which was sensitive to EB2023 *in vitro* (AML06) and a biallelic *TP53*-mutated sample which was insensitive to EB2023 *in vitro* (AML13) and selected subtherapeutic ven/aza doses known to induce minimal response in these same PDX models.^43^ As vehicle mice developed systemic leukemia at 8 weeks post-transplant, the experiment was terminated to measure the degree of AML infiltration in all animals. Near amelioration of tumor burden within the bone marrow was observed in both models, even at a dose where little monotherapy activity was observed (Fig. 4h, Extended Data Fig. 6j). This, again, highlights that the synergistic efficacy of EB2023 occurs at doses well below those needed to induce single agent responses.

The consistent anti-leukemic responses observed *in vitro* and *in vivo* provide preclinical rationale to evaluate EB2023 in clinical studies, but given recent on-target toxicities seen with OXPHOS inhibitors in clinical trials, we sought to further investigate how EB2023 effects metabolism and mitochondria in healthy tissue to better understand how chemically targeting the F_1_ subunit of ATP synthase could be best leveraged to define therapeutic index and treat human disease.

### EB2023 has unique pharmacokinetic properties and delivers a short duration energetic stress

In formal pharmacokinetic studies, C57/B6 mice were dosed via IV, IP and PO routes. EB2023 was quantifiable up to 2 hours after administration IV/IP, with 83.7% bioavailability and short plasma t_1/2_ of approximately 30 minutes after IV administration (Fig. 5a, Supplemental Table 2). This is a notable contrast to the reported T_1/2_ of I759 (16h).^15^ The phosphorylation of AMPK has been noted to be a reliable and dose-dependent biomarker of energetic stress from OXPHOS inhibitors in AML cells,^44^ and, as expected, this was the case as with EB2023 (Extended Data Fig. 5b). As it is known that AMPK becomes dephosphorylated within minutes of energy balance restoration,^45^ we postulated that dynamic monitoring of AMPK phosphorylation state could reveal differences in the energetic stress experienced by tissues based on duration of drug activity. To assess OXPHOS inhibitor-induced energetic stress *in vivo*, tissue AMPK phosphorylation was used as a pharmacodynamic surrogate of cellular ATP depletion (Fig. 5b). Phosphorylation of AMPK at Thr172 is a rapid and sensitive indicator of reductions in the ATP/AMP, making it well-suited for time-resolved assessment of drug activity in tissues.^44^ While we detected increased AMPK phosphorylation in mouse myocardium 2 hours after a single dose of 0.1 mg/kg EB2023, this was not the case in other tissues measured (Extended Data Fig. 7a). Importantly, the AMPK phosphorylation in mouse myocardium was transient; rising by 2 hours after a single EB2023 dose before decreasing and returning to baseline levels by 24 hours (Fig. 5c). In NSGS mice transplanted with MV-4-11 cells, serial phospho-AMPK measurements of AML cells from the bone marrow showed time-dependent decreases in AMPK phosphorylation by 24 hours with EB2023 to a greater extent than with I759 or oligomycin (Fig. 5d, Extended Data Fig. 7b), consistent with our hypothesis that EB2023 delivers an energetic stress to tissues for a shorter duration than either of the other OXPHOS inhibitors at doses that offer anti-AML activity in these same models. We propose a model of OXPHOS inhibitor toxicity (Fig. 5e) in which both C_max_ and the duration of metabolic stress drives on-target toxicity. With this model in mind, we sought to further clarify whether daily dosing of EB2023 prevented on-target toxicity seen with prior OXPHOS inhibitors.

**Figure 5.**
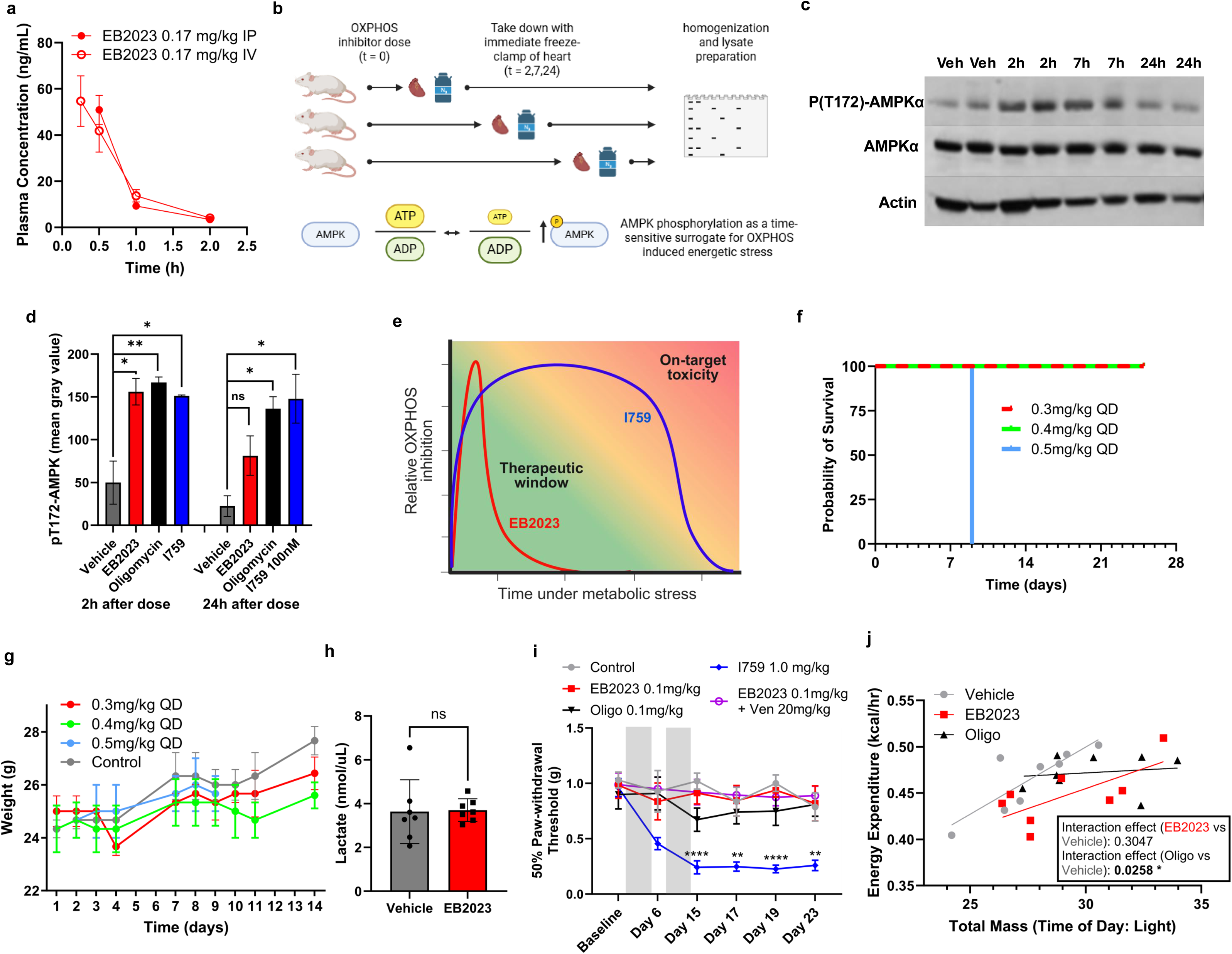
EB2023 avoids on-target toxicity on healthy tissues in mouse models and has a therapeutic window *in vivo*. **(a)** Pharmacokinetic studies of a single dose of EB2023 administered via IP or IV to C57BL/6 mice (n=3 per group) showing concentration of EB2023 as measured from serially sampled plasma by LC-MS. **(b)** Experimental schematic illustrating serial tissue harvest and analysis of AMPK phosphorylation following OXPHOS inhibitor dosing in mice. Created with BioRender.com. **(c)** Immunoblot analysis of phospho-AMPK (Thr172), total AMPKα, and actin in mouse myocardium harvested at 0, 2, 7, and 24 hours following a single dose of vehicle or EB2023 (0.1 mg/kg; n = 2 per timepoint). **(d)** Densitometric quantification of phospho-AMPK (Thr172) normalized to actin at 2 and 24 hours post-treatment in mice dosed with vehicle, EB2023, oligomycin, or I759 (n = 2–3 per group). Data shown as mean ± SEM. Comparisons performed by Welch’s unpaired t-test. **(e)** Schematic illustrating pharmacokinetic hypothesis for the differential therapeutic window of EB2023 versus I759, depicting time-dependent OXPHOS suppression relative to the therapeutic window and the time-dependent on-target toxicity threshold. Created with BioRender.com. **(f)** Kaplan-Meier survival curves for CD1 mice treated with EB2023 at 0.3, 0.4, or 0.5 mg/kg once daily (n= 6 per group; QD, 5 days on/2 days off). **(g)** Body weights of CD1 mice (n = 6 per group) treated with EB2023 at 0.3, 0.4, or 0.5 mg/kg once daily (5 days on/2 days off) for 2 weeks. Data shown as mean ± SEM. **(h)** Plasma lactate levels (nmol/µL) in CD1 mice (n = 7 per group) treated with vehicle or EB2023 (0.1 mg/kg, once daily, 5 days on/2 days off for 2 weeks), measured 18 hours after the final dose. Comparison performed by Welch’s unpaired t-test. **(i)** 50% paw withdrawal threshold (Von Frey mechanical allodynia testing) in CD1 mice (n = 5 per group) treated with EB2023 (0.1 mg/kg or 0.25 mg/kg, once daily) or I759 (1.0 mg/kg, once daily), venetoclax (20 mg/kg, once daily) or EB2023 and ven, all on a 5 days on/2 days off schedule. Comparisons performed by two-way ANOVA with Tukey’s post hoc test. **(j)** Correlation between total body mass and energy expenditure (kcal/hr) during the light phase in CD1 mice (n = 8 per group) treated with vehicle, EB2023, or oligomycin daily for 2 weeks using the Promethion Core Metabolic System (Sable Systems). Interaction effects between treatment and body mass on energy expenditure were assessed by ANCOVA; interaction p-values for EB2023 vs. vehicle and oligomycin vs. vehicle are shown. All bar and dot plots represent mean ± SEM. ns = not significant, *p<0.05, **p<0.01, ***p<0.001, ****p<0.0001.

### EB2023 does not induce evidence of on-target toxicity in mouse models at effective anti-AML doses

There were no clinically significant differences in blood count measurements of healthy CD1 mice treated with EB2023 for 2 weeks (Extended Data Fig. 7c). Consistent with this finding, EB2023 had a cytostatic but not cytotoxic effect on healthy CD34+ cells *in vitro*, whereas it decreased viability in MV-4-11 cells at the same timepoint (Extended Data Fig. 7d-e). Healthy CD34+ cells survived exposure to EB2023 despite similar degrees of activation of the ISR and OPA1 proteolysis (Extended Data Fig. 7f), suggesting healthy CD34+ cells may have greater mitochondrial stress adaptation capabilities.

In maximal tolerated dose testing, deaths were observed after 2 weeks of daily dosing at 0.5 mg/kg of EB2023 in a CD1 mouse model, with no deaths observed at lower doses (Fig. 5f). There were no observed decreases in mouse weight over 2 weeks with up to 0.4 mg/kg dosing (Fig. 5g) or in the triplet combination of EB2023/ven/aza (Extended Data Fig. 7g). Importantly, EB2023 did not cause an increase in serum lactate in CD1 mice (Fig. 5h), as was previously seen with I759 in mouse and human testing.^15^ Analogously, no evidence of peripheral neuropathy was observed with EB2023 daily dosing as measured by neurobehavioral adhesive removal or paw withdrawal testing, whereas the lowest reported effective dose of I759 (1.0 mg/kg) led to a significant reduction in the paw withdrawal threshold, a validated murine test for drug-induced neuropathy.^15,46^ Importantly, the addition of venetoclax did not potentiate peripheral neuropathy (Fig. 5i, Extended Data Fig. 7h).

To better understand healthy cell metabolism and how it is affected by ATP synthase inhibitors, we measured energetic and metabolic state using the Promethion Core Metabolic System. After dosing CD1 mice with oligomycin, EB2023, or vehicle control daily for 2 weeks, mouse energy expenditure, caloric intake, cage activity, and weights were measured. There were no statistically significant differences in weight loss, O2 consumption, CO2 production, food intake or energy expenditure between either ATP synthase inhibitor compared to vehicle (Supplementary Fig. 2a-f). While both ATP synthase inhibitors caused mild decreases in cage activity (Supplementary Fig. 2g-h), a dissociation of mouse lean body mass and energy expenditure was observed with oligomycin but not EB2023, evident as a loss of the expected positive correlation between lean body mass and energy expenditure (Fig. 5j). Such effects are observed with uncoupling weight loss compounds such as 2,4-dinitrophenol, and this effect is implicated in cardiotoxicity in humans, potentially mimicking the well-described cardiotoxicity of oligomycin observed in mouse models.^18,47^

## Discussion

For several decades, targeting cancer cell respiration directly has been stymied by a failure to demonstrate an acceptable therapeutic index with selective, potent inhibitors.^15–16,24,48^ Thus, any potential to address this vulnerability would require a nuanced approach to ETC disruption. Here, we conduct a comprehensive comparative characterization of the inhibition of processes bracketing OXPHOS and illustrate how F₁-selective ATP synthase inhibition is divergent from other strategies for OXPHOS inhibition.

While CRISPR-Cas9 depletion of ATP synthase subunits showed pan-essential effects across cancer lineages, chemical inhibition of this same enzyme demonstrated preferential activity against hematopoietic malignancies in our data mining efforts, implying that the pharmacology of enzyme inhibition, rather than complete loss of function by genetic depletion, could be a critical determinant of therapeutic index in OXPHOS inhibitor studies. Our finding that EB2023 maintains anti-AML efficacy while demonstrating improved tolerability compared to oligomycin and I759 supports this conclusion and is consistent with observations that the interplay between node of ETC inhibition and the temporal dynamics of ATP depletion are essential factors in evaluating OXPHOS inhibitors with a suitable therapeutic window.

F_1_ inhibition delivers three distinct advantages over other OXPHOS inhibitors. First, by targeting ATP synthase and permitting continued function of complexes I-IV (i.e. the respirasome) and the F_O_ subunit at therapeutic doses, EB2023 causes energetic stress (ATP depletion) while maintaining accompanying redox balance (NAD^+^/NADH preservation). I759 rapidly depletes the NAD^+^/NADH ratio and causes mitochondrial ROS accumulation due to complex I disruption, whereas EB2023 preserves NAD⁺/NADH ratios and does not increase ROS. Preservation of respirasome function temporarily permits continued NAD^+^-fueled TCA cycling and allows continued synthesis of critical nucleotide and biosynthetic precursors, attenuating the compensatory upregulation of glycolysis. While EB2023 had cytostatic effects on healthy CD34^+^ hematopoietic cells, viability was unaffected and lactate levels did not rise in mice treated daily with EB2023. The clinical significance is highlighted by recent findings that complex I inhibitor toxicity, particularly neurotoxicity, stems primarily from NADPH depletion rather than ATP depletion and the fact that glial cells are poorly equipped to handle this oxidative stress.^49^ Our observation that F_1_ inhibition at effective doses did not lead to any peripheral neuropathy, which was a primary limitation of I759, is a finding consistent with this mechanism and suggests that preserving NAD⁺/NADH redox balance while disrupting ATP synthesis may improve the therapeutic index of OXPHOS inhibitors.

Second, F_1_ inhibition with EB2023 delivers intermittent energetic stress *in vivo* that largely spares normal tissues from the effects of prolonged energy starvation; a hypothesis we tested with a novel method for surveying drug-induced energetic stress in model organisms by serial AMPK phosphorylation measurements. Phospho-AMPK measurement is a well-established rapid and dynamic reporter of AMP/ATP ratios,^45^ so we measured pAMPK in mouse myocardium and bone marrow from AML CDX and revealed that EB2023-induced energetic stress resolved while AMPK remained phosphorylated after I759 or oligomycin treatment for at least 24 hours. The transient nature of this energetic insult may explain why EB2023, unlike oligomycin, did not cause systemic metabolic uncoupling in mouse calorimetry experiments. These findings herald a greater appreciation of the temporal-dynamic effects in evaluation of OXPHOS inhibitors.

By comparing ATP synthase inhibitors in both a time dependent and dose dependent fashion, we also unveiled an unexpected divergence in Seahorse phenotypes from nanomolar to micromolar concentrations. At nanomolar concentrations, we observed continued oxygen consumption for as much as 4 hours after dosing either of the ATP synthase inhibitors. This is consistent with continued electron flux across complexes I-IV and proton flow across the inner mitochondrial membrane, without productive ATP synthesis at the inhibited F_1_ or F_o_ heads. However, at 20-fold higher concentrations (2 µM), both compounds produced immediate OCR suppression and glycolytic compensation, suggesting that supra-pharmacological glycomacrolide loading produces a qualitatively distinct effect, potentially reflecting multi-site occupancy or physical deformation of the F_o_F_1_ complex that impairs F_o_ proton conductance in addition to F_1_ catalytic activity. Despite decades of divergent reports in the literature with various oligomycin doses, these findings define two pharmacologically distinct concentrations for macrolide ATP synthase inhibitors, illustrating nuance between supra-pharmacological inhibition in metabolic assays and effective anti-cancer nanomolar concentrations *in vivo*.

Third, ATP synthase inhibition induced cell death more consistently across AML models than complex I inhibition, despite comparable ATP depletion and disruption of respiration. Initial studies of these F_1_ inhibitors noted that potent ATPase inhibition was insufficient to explain their antiproliferative activity, suggesting secondary biological effects beyond ATP depletion.^50^ We propose that on-target, downstream mitochondrial stresses induced by EB2023 contribute to its potent anti-AML effect and that these effects are unique to ATP synthase inhibitors. The integrated stress response (ISR), known to be activated via the OMA1-DELE1-HRI axis in response to ATP synthase inhibition,^51^ plays a critical role in the anti-AML activity seen with F_1_ inhibition. Understanding the distinct mechanisms by which OXPHOS inhibitors trigger mitochondrial stress responses and how different cell types process these stresses is critical.

Our study identifies OPA1-mediated mitochondrial structural remodeling as another unique property of ATP synthase inhibition. Proteolytic cleavage of l-OPA1 to s-OPA1 results in unopposed fission and cristae remodeling, which, if sustained, leads to mitochondrial integrity loss and caspase-independent cell death.^36,52^ Prior work in HeLa cells showed that only ATP synthase inhibition, and not chemical inhibition of complex I or III, triggers this change in OPA1,^53^ consistent with our findings comparing EB2023 and I759. The exact mechanism by which ATP synthase disruption triggers OMA1 proteolytic cleavage of OPA1 is incompletely understood but it is thought to be distinct from mitochondrial membrane potential loss and OMA1 has been hypothesized to sense ATP synthase uncoupling or stalling via interactions with Bax and Bak.^54–56^ Our work suggests that ATP synthase functional state interacts with OMA1 and triggers proteolytic activity which is divorced from energetic stress and mitochondrial membrane potential but the precise molecular mechanism by which ATP synthase stalling is transduced to OMA1 activation requires further investigation.

Importantly, recent reports suggest that OPA1 perturbations can overcome acquired venetoclax resistance and mitochondrial structural disruption has also been shown to sensitize AML cells to BCL2 inhibition.^34^ *De novo* or acquired venetoclax resistance remains a key challenge in the treatment of AML. Multiple metabolism-based mechanisms of venetoclax resistance have been demonstrated including fatty acid oxidation, increased OXPHOS, and mitochondrial structure changes, all of which are theoretically circumvented by terminal OXPHOS inhibition at ATP synthase.^35,57,58^ Notably, F_1_/BCl2 inhibition synergy occurs at relatively benign doses of both single agents and does not require maximal suppression of cellular respiration. This suggests that BCL2i synergistic benefit of F_1_ inhibition can be achieved without risking deleterious rewiring of healthy tissue metabolism. *RAS*-mutated AML, which is frequently of myelomonocytic or monocytic morphology, is particularly challenging to manage in the clinic, and is typically resistant to BCL2 inhibition.^59,60^ Thus, the capacity of F_1_ inhibition to sensitize *NRAS* and *TP53* mutated cells to BCL2 inhibition represents a translationally relevant advance.

Our systematic comparison establishes F_1_-selective ATP synthase inhibition as a promising AML therapeutic approach. EB2023, a F_1_ ATP synthase inhibitor, delivers a redox-balanced energetic stress, and it has unique pharmacokinetics with an acceptable therapeutic window to explore clinically in AML. Taken together, we have shown how F_1_ ATP synthase inhibition affects its anti-AML activity from mechanisms, at least in part, not fully related to its ability to deplete ATP, and that these pathways also drive BCL2 dependence and thus susceptibility to venetoclax. These findings provide both a path forward for the clinical development of small molecules that selectively inhibit F_1_ ATP synthase and establish novel principles for consideration of next-generation mitochondrially-targeted cancer therapeutics.

## Methods

### CRISPR gene dependency, gene expression and drug lineage enrichment analyses

Gene set lists for four metabolic processes were obtained from Harmonizome 3.0, using the GO Biological Process Annotations 2025 dataset. The analyzed gene sets were glycolytic process (GO:0006096; 29 genes), glutamine metabolic process (GO:0006541; 13 genes), fatty acid oxidation (GO:0019395; 52 genes), and oxidative phosphorylation (GO:0006119; 61 genes). Exact gene symbols used for each gene set are provided in Source Data. Genes present in the Harmonizome gene sets but absent from the DepMap CRISPR gene effect matrix were recorded as missing and excluded from mean calculations and statistical comparisons. Electron transport chain complex gene groupings (Complexes I–V) and ATP synthase subunit groupings (F1 subunit, FO subunit, and peripheral stalk) were defined using HUGO Gene Nomenclature Committee (HGNC) gene group classifications (genenames.org). Mean Chronos dependency scores were compared between selected gene groups (Complex I vs Complex V; F1 vs FO subunit) using unpaired Mann-Whitney U tests in GraphPad Prism for both the entire cancer cell line data set (n=1183) and AML models (n=30) as defined by models annotated in DepMap as OncotreePrimaryDisease = Acute Myeloid Leukemia. Drug sensitivity data for oligomycin (area under the curve, AUC), venetoclax (AUC), and IACS-010759 (log₂ fold change) were downloaded from the DepMap portal (depmap.org), sourced from the CTD² drug sensitivity dataset and PRISM Repurposing Public 24Q2 dataset, respectively. Lineage annotations were assigned using DepMap’s built-in cell line classifications. Differences in drug sensitivity between hematopoietic sub lineages and all other cancer cell lines were assessed by Mann-Whitney U test in GraphPad Prism. PDK1 gene expression data were obtained from the DepMap Public 25Q2 OmicsExpressionProteinCodingGenesExpectedCountsProfile dataset, which provides log-transformed TPM (transcripts per million) values derived from unstranded RNA-seq for protein-coding genes. AML cell lines (n = 10) were ranked by EB2023 sensitivity (AUC from CellTiter-Glo dose-response assay) and divided into the five most and five least sensitive lines. PDK1 expression was compared between groups by Mann-Whitney U test in GraphPad Prism.

### NCI60 cancer cell line growth screen

Anti-cancer activity of apoptolidin A, the parent compound of the F_1_ ATP synthase-targeting glycomacrolide family, was assessed across the NCI-60 cancer cell line panel by the National Cancer Institute Developmental Therapeutics Program (NCI DTP) at a single concentration of 1000 nM under standard NCI-60 screening conditions.

### Patient samples

Primary AML patient samples were provided by the Vanderbilt-Ingram Cancer Center Hematopoietic Malignancies Repository and experiments were conducted in accordance with the tenets of the Declaration of Helsinki and approved by the Vanderbilt University Medical Center Institutional Review Board. Written informed consent was obtained from all patients. Samples were obtained either from leukapheresis procedures or bone marrow aspirates, as detailed in Supplementary Table 1. Mononuclear cells were isolated by density gradient centrifugation and used fresh or cryopreserved until use. Human CD34+ primary umbilical cord blood samples were purchased from Stem Cell Technologies.

### Next-generation sequencing

For next-generation sequencing (NGS), AML patient bone marrow aspirates or leukapheresis samples were obtained, and DNA was isolated using a DNA Midi Kit (Qiagen, Hilden, Germany) for NGS in a panel of commonly mutated regions of myeloid neoplasia-associated genes. The analytic targets included in the OnkoSight NGS Myeloid Malignancies gene panel (Illumina, San Diego, CA) include exonic regions across each of the following genes: SRSF2, U2AF1, TET2, IDH2, DNMT3A, RUNX1, TP53, BCOR, BCORL1, ETV6, NPM1, GATA2, WT1, ASXL1, EZH2, JAK2, FLT3, FBXW7, CBL, KRAS, NRAS, SETBP1, ABL1, CSF3R, PTEN, PTPN11, ZRSR2, PHF6, MYD88, IDH1, HRAS, CALR, BRAF, and CDKN2A. The panel of validated genes consisted of therapeutic markers as well as genes with diagnostic and prognostic utility in myeloid and other hematologic tumors. Sample-level molecular annotations are provided in Supplementary Table 1.

### Isolation of EB2023 (ammocidin A) and apoptolidin A

Identical methods were used as described previously.^19^ Apoptolidin A was obtained by cultivation of wild-type *Nocardiopsis* sp. FU-40 and EB2023 (ammocidin A) was obtained by cultivation of *Saccharothrix* sp. AJ9571 provided by Ajinomoto Co. Each organism was plated on Bennett’s agar (0.1% yeast extract, 0.1% beef extract, 0.2% N-Z Amine A, 1.0% dextrose, 2.0% agar, pH 7.0) and incubated at 30 °C for 3–7 days until sporulation. FU-40 Δ*ApoGT2* was grown on plates containing Bennet’s media with the addition of apramycin, 80 μg ml^−1^. The seed culture was initiated using spores scraped from the solid culture into 250-ml Erlenmeyer flasks containing 50 ml of seed medium (1.0% soluble starch, 1.0% molasses (Plantation Blackstrap, unsulfured), 1.0% peptone, 1.0% beef extract, pH 7.0) and incubated for 7 days at 30 °C while shaking at 220 rpm. Production cultures were carried out in multiple 250-ml Erlenmeyer flasks containing 50 ml of production media (2.0% glycerol, 1.0% molasses, 0.5% casamino acids, 0.1% peptone, 0.4% calcium carbonate, pH 7.2) and incubated at 30 °C for 7 days while shaking at 220 r.p.m. Mycelia were then separated from the culture broth by centrifugation at 3,000*g* for 30 min. The culture broth was extracted with 3× 1 volume of ethyl acetate and the combined organic layers were washed with brine, dried with Na_2_SO_4_ and concentrated in vacuo. The crude extracts were then subjected to chromatography with LH-20 resin using methanol as the mobile phase and the glycomacrolide-containing fractions were identified by thin layer chromatography and pooled. The LH-20 fractions were then subjected to reverse-phase HPLC using a Waters XBridge Prep C_18_ 19 × 150 mm column with a 20-min gradient from 70% A/30% B to 20% A/80% B (buffer A: 95% water, 5% acetonitrile, 10 mM ammonium acetate; buffer B: 5% water, 95% acetonitrile, 10 mM ammonium acetate)

### Cell lines

AML cell lines MV-4-11, THP-1, and U-937 were purchased from the ATCC. MOLM-13, OCI-AML-3, KG-1, NB-4, SET-2, OCI-AML-5, and HEL cell lines were purchased from Deutsche Sammlung von Mikroorganismen und Zellkulturen (DSMZ). ATCC and DSMZ cell bank cell lines are authenticated by short tandem repeat profiling and cytochrome b oxidase gene sequencing. Cultured cells were split every 3–4 days and maintained in exponential growth phase. Cell lines were tested for Mycoplasma contamination using the Mycoplasma PCR Detection Kit (abcam; ab289834). Cells were used within 30 passages from thawing. MV-4-11 cells were grown in IMDM; OCI-AML-3 in alpha-MEM; all other cell lines in RPMI-1640. All media were supplemented with 10% FBS and 100 U/mL penicillin and 100 µg/mL streptomycin. Cells were maintained at 37°C in a 5% CO₂ incubator.

### Cell viability assays

#### CellTiter-Glo

Compounds were diluted in DMSO (final DMSO concentration 0.2%) and dispensed into 384-well plates using the Echo 650 acoustic liquid handler (Labcyte) with EB2023, IACS-010759, and oligomycin at concentrations from 1 nM to 10 µM, and venetoclax from 31 nM to 1 µM. Following compound addition, cells were plated at 2,000–8,000 cells per well in appropriate media supplemented with 10% FBS and incubated at 37°C, 5% CO₂. Plates were incubated for 48 or 72 hours and cell viability was measured using the CellTiter-Glo reagent (Promega) per manufacturer’s protocol on a Cytation 1 multimode reader (BioTek/Agilent). Percent viability was defined as relative luminescence units (RLU) normalized to DMSO-treated controls. Ten-point dose-response curves were fit by nonlinear regression using a four-parameter logistic model in GraphPad Prism, and IC_50_ values were determined from the resulting fits The Echo 650 Series Acoustic Liquid Handler is housed and managed within the Vanderbilt High-Throughput Screening Core Facility, an institutionally supported core

#### CellTiter-Fluor

Cell viability was orthogonally assessed using the CellTiter-Fluor^TM^ Cell Viability Assay (Promega Corp.), which measures viable cell protease activity as a fluorescent readout independent of cellular ATP content. Assay conditions, cell plating, compound dilutions, and incubation parameters were identical to those described for CellTiter-Glo above. Fluorescence was measured per manufacturer’s protocol and percent viability was calculated relative to DMSO-treated controls.

#### Trypan exclusion

Cells were treated with indicated compounds under standard culture conditions for 24–72 hours. At each timepoint, 20 µL aliquots were collected from each well (n = 4 replicates per condition) and mixed 1:1 with trypan blue stain for 15 seconds. Viability and total cell counts were measured on a Countess 3 automated cell counter (Thermo Fisher Scientific). Percent viability was expressed relative to DMSO-treated controls.

#### Caspase 3/7 activity assessment

Caspase 3/7 activity was assessed using the Caspase-Glo^TM^ 3/7 Assay System (Promega Corp.) according to the manufacturer’s protocol. Cells were treated with indicated compounds under standard culture conditions for 24 hours, after which Caspase-Glo 3/7 reagent was added directly to culture wells in a 1:1 volume ratio and incubated at room temperature for 30 minutes. Luminescence was measured on a Cytation 1 multimode plate reader (BioTek/Agilent). Caspase 3/7 activity was expressed as relative light units (RLU) normalized to DMSO-treated controls.

### Flow cytometry

For flow cytometry, red blood cells were lysed with EL Buffer on ice (Qiagen), with remaining cells washed and resuspended in 1× PBS with 1% BSA and stained for 15 minutes with the following antibodies: human CD45-APC (Clone 2D1; BioLegend), human CD33-PE-Cy7 (Clone P67.6; BioLegend), murine CD45-PE (Clone 30-F11; BioLegend), and DAPI (BioLegend). For cell cycling using Ki67 and DAPI, cells were fixed/permeabilized with Cytofix Kit (Becton Dickinson) and incubated overnight with APC-Ki67 antibody (BioLegend) and stained with DAPI 5 minutes prior to flow cytometric analysis. Cells were washed and submitted for flow cytometric analysis using a three-laser LSRII (Becton Dickinson). For annexin/PI staining, an annexin V apoptosis kit was used as per manufacturer’s instructions (BD Pharmingen). For patient samples, mononuclear cells (MNCs) were subjected to 24 hours of drug treatment and stained for flow cytometry. Primary AML blast cells were gated as CD45^lo-mid^/CD33^hi^/SSC-A^lo^.

### Extracellular flux analyses (Seahorse)

OCR and ECAR were determined using commercial Mito Stress test kit (Agilent, catalog No. 10301) and Glycolysis Stress Test kit (Agilent, catalog No. 103020) per manufacturer’s protocol and experiments were carried out on Seahorse XF96 bioanalyzer (Agilent). MV-4-11, MOLM-13, and THP-1, cells were treated for 24 hours with 100 or 20000 nmol/L EB2023, oligomycin or IACS-010759 for OCR and ECAR analysis. After treatment, cells were collected and washed in PBS. Cell viability was determined via Trypan Blue on Bio-Rad TC20 automated cell counter. Each well was loaded with 7.5 × 10⁴ live cells. Cells were spun onto XF96 Cell-Tak (BD Biosciences, catalog No. 354240) coated plates and rested in Seahorse XF RPMI1640 media supplemented with glutamine, sodium pyruvate, and glucose. Extracellular acidification rate (ECAR) and oxygen consumption rate (OCR) values were normalized to number of live cells per well measured immediately before plating by trypan blue exclusion on Countess 3 automated cell counter. For the continuous assessments after OXPHOS inhibitor dosing (Fig. 2d and Extended Data Fig. 2c), cells were plated as above and the Seahorse XF96 bioanalyzer was loaded with each OXPHOS inhibitor. OCR and ECAR of untreated cells were measured every 5 minute to establish a baseline, then drug was auto-injected after 30 minutes and measurement frequency was increased to every 2 minutes for a 240-minute duration of measurements.

### LC–MS for targeted metabolomics and stable-isotope tracing in AML cell lines

MV-4-11 and Molm-13 were cultured in RPMI-1640 media supplemented with 10% dialyzed FBS, 2 mM [^13^C_5_,^15^N_2_]-glutamine (Cambridge Laboratories) and 2 g/L unlabeled glucose and treated with DMSO control, EB2023 100nM, oligomycin 100nM, or IACS-010759. After 24 hours, cells were centrifuged, washed twice with ice-cold PBS and flash-frozen in liquid nitrogen. Polar metabolites were extracted for LC-MS analysis using a modified Bligh-Dyer procedure.^61^ Polar fraction was acquired with an Orbitrap IQ-X Tribrid mass spectrometer coupled with the ultra-high-pressure liquid chromatography (UHPLC) Vanquish pump with climate controlled (4 °C) autosampler (Thermo Scientific, Waltham, MA, USA). To optimize polar metabolites analysis, two columns were used as previously described.^62,63^ Peak picking and alignment were performed with SIEVE 2.2 software (Thermo Scientific, Waltham, MA, USA) using a mass tolerance of 5 ppm and retention time (RT) tolerance of 0.25 minutes. Peaks were then scaled according to probabilistic quotient normalization.^64^ Metabolites annotation was conducted by matching mass to charge ratio (m/z) and RT to a library of standards generated in house that include IROA 300 MS Metabolite Library of Standards (IROA Technologies) or by putatively by matching m/z to the KEGG database using an in-house MATLAB script.^63^ Isotope natural abundance correction was performed using AccuCor2.^65^

### Reactive oxygen species/tetramethylrhodamine methyl ester for flow cytometry

Mitochondrial membrane potential was assessed using the MitoProbe^TM^ TMRM Assay Kit for Flow Cytometry (Invitrogen) per manufacturer’s protocol at a non-quenching TMRM concentration of 20 nM. Mitochondrial ROS was assessed using MitoSOX^TM^ Mitochondrial Superoxide Indicator (Invitrogen) per manufacturer’s protocol. Cells were pretreated with the indicated OXPHOS inhibitor and duration prior to dye incubation directly in cell culture media. CCCP served as a positive depolarization control for TMRM experiments. Cells were incubated with dye for 30 minutes at 37°C protected from light, washed twice, and analyzed on a three-laser LSRII flow cytometer (BD Biosciences).

### mRNA-seq library preparation and sequencing

Total RNA was extracted from MV-4-11 and THP-1 cells treated with EB2023, oligomycin, or IACS-010759 (100 nM each) or DMSO control for 2 or 24 hours using a standard RNA isolation protocol (Rneasy Mini Kit, Qiagen). RNA quality and quantity were assessed prior to submission. mRNA library preparation and sequencing were performed by Genewiz (Azenta Life Sciences). Briefly, mRNA was enriched from total RNA by poly(A) selection, fragmented, and converted to cDNA. Following cDNA synthesis, end repair, A-tailing, and adapter ligation were performed, and libraries were amplified by PCR. Sequencing was performed on an Illumina NovaSeq platform with a paired-end 2 × 150 bp configuration at a minimum sequencing depth of 20–30 million read pairs per sample, with a guaranteed data quality of ≥85% bases at Q30 or higher.

### RNA-seq analysis

Adapter sequences were trimmed from raw reads using Cutadapt (v4.5). Quality control was performed on both raw and adapter-trimmed reads using FastQC (v0.12.1; www.bioinformatics.babraham.ac.uk/projects/fastqc). Trimmed reads were aligned to the human reference genome GRCh38.p13 using STAR (v2.7.11a), with Gencode v38 gene annotations provided to improve mapping accuracy. Read counts per gene were quantified using featureCounts (v2.0.6). Differentially expressed genes were identified using DESeq2 (v1.42.1); genes with absolute log₂ fold change > 0.5 and FDR-adjusted p-value ≤ 0.05 were considered statistically significant. Volcano plots were generated in Prism, with significance thresholds set at |log₂ fold change| > 0.5 and −log₁₀(p-value) > 2. Gene set enrichment analysis was performed using WebGestalt (WebGestaltR v0.4.6) against the Reactome pathway database (v2022.1.Hs), with genes ranked by the DESeq2 Wald statistic, a minimum of 5 and maximum of 2,000 genes per category, 1,000 permutations, and redundancy removal by weighted set cover. Results were visualized as normalized enrichment score (NES) heatmaps generated using GraphPad Prism.

### Quantitative immunoblot

Cells were grown under indicated treatment conditions. Total protein lysates were extracted from 6.0 × 10⁶ cells per condition in Laemmli sample buffer (Bio-Rad), sonicated, and boiled at 95°C for 10 minutes. Samples were resolved by 10% SDS-PAGE and transferred to PVDF membranes. Immunoblot analysis was performed according to standard protocol with the following antibodies from Cell Signaling Technology: AMPKα (2532), phospho-AMPKα (Thr172; 2531), ATF4 (D4B8; 11815), ATF6 (D4Z8V; 65880), CHOP (L63F7; 2895), phospho-eIF2α (Ser51; 9721), eIF2α (9722), OPA1 (6589), MCL1 (94296); and actin (Sigma-Aldrich). Membranes were imaged and band densitometry was performed using ImageJ. Protein abundance was expressed as the ratio of band intensity of the protein of interest relative to actin loading control.

### Transmission electron microscopy analysis of AML cells

For transmission electron microscopy (TEM), samples were obtained and placed into 2.5 % glutaraldehyde in 1x PBS and postfixed sequentially in 1 % tannic acid, 1 % OsO4, and stained with 1 % uranyl acetate. Samples were then dehydrated in a graded ethanol series and infiltrated with EMbed-812 resin (Electron Microscopy Sciences, Hatfield, PA, USA) using propylene oxide as the transition solvent. The resin was polymerized at 60 °C for 48 h. Thin sections were taken at a nominal thickness of 70 nm and collected on 200 mesh Ni grids. Grids were stained with 1 % phosphotungstic acid followed by 1 % UA and lead citrate. Imaging was carried out with a Tecnai T12 operating at 100 kV equipped on an AMT nanosprint CMOS camera using SerialEM acquisition software.

For morphometric analysis of mitochondrial ultrastructure, ImageJ software was used to quantify mitochondrial lucency, roundness, circularity, aspect ratio, and cross-sectional area from TEM images. An unbiased, representative selection of cell images was compiled by the imaging core without prior knowledge of treatment condition, and 200 individual mitochondria per treatment condition were identified and manually traced as regions of interest (ROI). Mitochondrial electron lucency was calculated as the difference in mean gray value between the extracellular space (background) and the mitochondrial matrix ROI, where higher values indicate a less electron-dense, more electron-lucent matrix. Mitochondrial roundness and circularity were calculated by ImageJ as 4π × area/perimeter² and 4π × area/perimeter², respectively, on a unitless scale from 0 to 1 where values approaching 1 indicate a more circular morphology. Aspect ratio was calculated as the ratio of the major to minor axis of the best-fit ellipse. ROI area was recorded in calibrated units as a measure of mitochondrial cross-sectional size.

### *In vivo* murine modeling

All animal experiments were conducted in accordance with guidelines approved by the IACUC at Vanderbilt University Medical Center (Nashville, TN). Female NSGS [NOD-scid IL2Rgnull3Tg (hSCF/hGM-CSF/hIL3)] mice, 6–8 weeks old were irradiated with 100 cGy microwave radiation. Twenty-four hours later, mice were transplanted intravenously with cells of interest. In the cell line–derived xenografts (CDX), 1 × 10^6^ MV-4-11 cells were transplanted via tail vein injections in each irradiated mouse. For patient sample–derived xenografts (PDX), 2 × 10^6^ AML MNC were transplanted via tail vein injections. Mice were randomized post cell injection into cages of five. Prior to treatment, peripheral microchimerism was documented at week 1 in CDX. For AML PDX, peripheral chimerism was established by 2 weeks. Mice showing no peripheral chimerism by 2 weeks in CDX, or 3 weeks in AML PDX, were considered engraftment failures and removed from the study. Upon establishing microchimerism, mice were treated with either venetoclax (VEN, Chemietek), azacytidine (AZA, Chemietek) or EB2023. Venetoclax was dissolved in polyethylene glycol and ethanol and diluted with Phosal 50 PG for gavage. EB2023 and AZA were dissolved in DMSO and diluted in buffered saline for intraperitoneal injection. Mice were treated with VEN and AZA (VEN 20 mg/kg 5 days [d] on 2 d off, AZA upfront Monday/Wednesday/Friday [M/W/F] 1.5 mg/kg), either with or without EB2023 (EB2023 0.1 mg/kg 5D on 2 D off). Peripheral blood was assessed weekly for human chimerism. Spleen/body ratio was calculated as organ weight (gram) per gram of body weight. Murine complete blood count was conducted on blood collected into EDTA tubes (Greiner Bio-One) and analyzed with a Hemavet (Drew Scientific) analysis system.

### Histology and IHC

Tissues were fixed in 4% paraformaldehyde for 48 hours and stored in 70% ethanol before being embedded in paraffin and sectioned at 5 μm. The bone tissue was decalcified prior to being embedded in paraffin. Sections were dewaxed in xylene and rehydrated in successive ethanol baths. Standard Mayer hematoxylin and eosin (H&E) staining was performed. Antigen retrieval using a standard pH 6 sodium citrate buffer (BioGenex) was performed and sections were stained with Monoclonal Mouse Anti-Human CD45 (Dako, M0701, dilution 1:200) using the M.O.M. Kit (Vector).

### BH3 Profiling

BH3 profiling was conducted as previously reported.^66^ In brief, THP-1 cells were pretreated with 300 nM EB2023 for 24h, permeabilized with digitonin and exposed to BH3 peptides (BIM, BID, PUMA, Bmf-Y, HRK, NOXA, MS-1, BAD, and FS-1; synthesized by New England Peptide), and BH3 mimetics S63845, ABT-199 (venetoclax) or ABT-263. Alameticin (Ala) was used as positive control leading to 100% cytochrome c release. Cells were exposed to BH3 peptides and mimetics at 25°C for 60 min before fixation in 4% PFA, neutralization, and staining with Hoechst 33342 (Thermo Fisher) and anti-cytochrome c antibody (Biolegend) and analyzed by flow cytometry. The raw median fluorescence intensity (MFI) data were converted to the cytochrome c release parameter as follows: Cytochrome C release = [1-(MFI_sample_-MFI_Ala)_/(MFI_DMSO_-MFI_Ala_)] and normalized to DMSO control cells.

### Sample preparation and LC-MS/MS pharmacokinetic analysis

Young, adult male C57BL/6 mice, weighing 20-25 g were randomly divided into 13 groups of N = 3 each and EB2023 was administered via IV, IP and PO routes at a dose of 0.170 mg/kg. The mice were euthanized at 0.25 (IV group only), 0.5, 1, 2, and 4 h, post-dose and blood samples were collected in heparinized tubes. Plasma was harvested by centrifuging the blood at 10000 rpm for 10 min and stored frozen at –80°C until bioanalysis.

A simple protein precipitation method was used for the extraction of EB2023 from plasma samples. A 20 µl plasma sample was treated with acetonitrile containing 20 ng/mL of verapamil as internal standard (I.S.) at a ratio of 1:3 and vortex-mixed for 5 minutes followed by filtration through 0.45 µm filter plates (Millipore Solvinert® plates) by centrifugation at 1500 rpm for 5 min. UPLC-MS/MS analysis was carried out using Waters Acquity Class I Plus UPLC coupled with a Waters Xevo TQ-XS triple quadrupole mass spectrometer (MS/MS). The chromatographic separation was achieved using Acquity UPLC BEH C18 column (2.1 mm x 50 mm, 1.7 µm) and the mobile phase consisted of 0.1% formic acid (A) – acetonitrile (B) with a gradient program of 90 % A held for 0.3 min, decreased to 70% reaching 1.0 min followed by decreasing to 50% in 0.5 min. Then %A was further decreased to 10% reaching 2.25 min, held until 3.0 min and sharply decreased back to the initial conditions by 3.1 min and maintained until 3.5 min. The column and autosampler temperatures were kept at 55 °C and 10 °C, respectively. The mobile phase was delivered at a flow rate of 0.35 mL/min and the injection volume was set to 4 μL. The MassLynx software version 4.2 was used for instrument control and TargetLynx for data analysis. The mass spectrometer was operated in positive ion mode and detection of the ions was performed in the multiple reaction monitoring (MRM) mode, monitoring transition of m/z 1139.6 precursor ion [M+H]+ to the m/z 209.1 product ion for EB2023 and m/z 455.3 precursor ion [M+H]+ to m/z 150.1 product ion for I.S. (verapamil). The cone voltage and collision energy were 20, 22 V for EB2023, and 28, 42 V for verapamil. Test samples were analyzed with freshly prepared calibration and quality control standards for a linearity range of 2-1000 ng/mL in plasma.

### Behavioral studies assessing treatment-induced peripheral neuropathy

To investigate the effects of treatment-induced peripheral neuropathy, male and female CD1 mice were randomized into groups (n = 5 mice per group) that received either EB2023 (0.1 or 0.25 mg/kg in normal saline, i.p.), IACS-010759 (1.0 mg/kg in 0.5% methyl cellulose, p.o.), oligomycin (0.1 mg/kg in normal saline, i.p.), venetoclax (20 mg/kg in 60% PHOSAL 50 PG + 40% volume of diluent combination of 75% PEG & 25% ethanol, p.o.) or vehicle (control). All treatments were administered once per day, 5 days on, 2 days off, followed by a second round of 5 days on.

Mechanical allodynia was quantified using von Frey-calibrated filaments at baseline (before treatment), day 6 (after the first cycle of doses), day 15, 17, 19 and day 23. The mechanical stimulus inducing 50% likelihood of withdrawal was determined using the ‘up-down’ calculation method as previously described.^67^ Briefly, mice were habituated on a wire-grid panel for 20 min before testing. A series of von Frey filaments (0.16, 0.40, 0.60, 1.00 and 1.40 g, Stoelting) was applied to the plantar surface of both hind paws. Rapid withdrawal or flinching of the hind paw was considered a positive response. In the absence of a response, the next greater force was applied.

Sensory function was assessed using the adhesive removal test as previously described.^67^ Briefly, a round adhesive patch (3/15” Teeny Touch-Spots, USA Scientific INC, USA) was placed on the plantar surface of the mouse hind paw. The animals were then released into a clean plexiglass box, and the latency to detect and respond to the adhesive stimulus was recorded. The response time was defined as the time from placement in the arena to the first attempt to contact, shake, or remove the adhesive from the paw. Each mouse was tested in 6 trials (3 trials for each hind paw), and the average response latency was used for analysis.

### Measurement of healthy organ tissue AMPK phosphorylation

Following a single dose of OXPHOS inhibitor administered by intraperitoneal injection, mice were sacrificed at 2, 7, or 24 hours post-dose, with an untreated cohort serving as baseline control. Immediately upon sacrifice, tissues were harvested and freeze-clamped using a liquid nitrogen-cooled hand clamp to arrest metabolic activity and preserve phosphorylation state at the moment of harvest. In an initial experiment assessing tissue distribution of energetic stress (Extended Data Fig. 7e), heart, liver, brain, spleen, and kidney were harvested at 2 hours post-dose. In subsequent time-course experiments (Fig. 5h–i), cardiac (h) and bone marrow (i) tissue was harvested at 0, 2, 7, and 24 hours post-dose.

Frozen tissue was processed over dry ice to prevent thawing. Tissue fragments were shaved with a sterile scalpel into fine particles and transferred to lysis buffer, then sonicated to achieve complete homogenization. Protein lysates were prepared and subjected to immunoblot analysis using identical conditions to those described for cell line western blotting above. Membranes were probed for phospho-AMPKα (Thr172, Cell Signaling 2531) and total AMPKα (Cell Signaling 2532), with actin (Sigma Aldrich) serving as loading control. Densitometric quantification was performed using ImageJ.

### Promethion analysis and plasma lactate quantification

Mice were individually placed in home-cages in a 12h light/dark cycle, temperature/humidity-controlled dedicated room located in the Vanderbilt MMPC (RRID: SCIR_021939). Cohorts of 8 mice were treated with either vehicle of DMSO in saline I.P., EB2023 0.1mg/kg I.P. daily (D1-5,8-12) or oligomycin 0.1mg/kg I.P. daily (D1-5,8-12) with calorimetry analysis taking place over the final 3 days of the two-week treatment course.

Energy expenditure measures were obtained by indirect calorimetry (Promethion, Sable Systems, Las Vegas, NV). The calorimetry system consists of home cages with bedding equipped with water bottles and food hoppers connected to load cells for food and water intake monitoring. All animals had *ad libitum* access to standard chow and water. The air within the cages is sampled through microperforated stainless steel sampling tubes that ensure uniform cage air sampling. Promethion utilizes a pull-mode, negative pressure system with an excurrent flow rate set at 2000 mL/min. Water vapor is continuously measured and its dilution effect on O_2_ and CO_2_ are mathematically compensated for in the analysis stream.^68^ O_2_ consumption and CO_2_ production are measured for each mouse every 5 mins for 30 seconds. Incurrent air reference values are determined every 4 cages. Respiratory quotient (RQ) is calculated as the ratio of CO_2_ production over O_2_ consumption. Energy expenditure is calculated using the Weir equation: EE (kcal/hr) = 60*(0.003941*VO_2_(mL/min) +0.001106*VCO_2_(mL/min)).^69^ Ambulatory activity was determined every second with XYZ beams. Data acquisition and processing were coordinated by MetaScreen and MacroInterpreter (Sable Systems). Calorimetry data were analyzed using CalR2. Mass-dependent variables (VO₂, VCO₂, energy expenditure, and food intake) were analyzed by ANCOVA with body mass as covariate; mass-independent variables (locomotor activity, respiratory quotient) were analyzed by ANOVA. Plasma lactate was measured from tail vein blood collected 18 hours after the final dose using a colorimetric lactate assay per manufacturer’s instructions.

### Statistical analysis

Unless otherwise noted, data were summarized using the mean (± SEM). Per group sample sizes are presented in figures and results reported from two independent experiments, unless stated otherwise. Pairwise comparisons were performed using Welch’s unpaired t-test or the nonparametric Mann-Whitney U test as indicated in figure legends. Growth curve data were analyzed by two-way repeated measures ANOVA with Dunnett’s post-hoc test versus DMSO control. Dose-response curves were fit by nonlinear regression using a four-parameter logistic model; Emax values (bottom plateau) were compared by extra sum-of-squares F-test. ¹³C isotope incorporation data were analyzed by two-way ANOVA with treatment and metabolite as factors, followed by Welch’s unpaired t-test with Holm-Bonferroni correction for post-hoc pairwise comparisons. Nucleotide abundance data were compared by Welch’s one-way ANOVA with Holm-Bonferroni corrected post-hoc Welch’s t-tests versus control. Primary patient sample viability data were analyzed by Friedman test comparing each treatment condition. Cell cycle phase distributions were compared by one-way ANOVA with Dunnett’s post-hoc test. Zero-interaction potency (ZIP) modeling was applied to the dose response matrices to assess drug synergy, represented by the ZIP score, which is the average combinatory effect of both drugs over the entire matrix of tested concentrations.^37^ Von Frey data were analyzed by repeated measures two-way ANOVA, with Tukey’s post hoc tests, and sticker removal data by one-way ANOVA with Tukey’s post hoc tests. Data were analyzed using Graph Pad Prism 6.0 for Windows (GraphPad Software, www.graphpad.com).

## Funding Acknowledgments

R35 GM154838 (EGT, AJM); R01 CA227064 (AKS, PMG); R01 CA226833 (BB); R01 CA231364-06 (to MK, AS, SC, ST); Conquer Cancer, The ASCO Foundation, T32 5T32CA217834-08 (MTV); P30 CA68485, R21 CA318444, Adventure Alle Fund and Beverly and George Rawlings Endowment (MTV, HER, MPA, YFA, MJ, AG, MRS).

The following scientific core services utilized in this work are supported by the NIH as noted below. The content is solely the responsibility of the authors and does not necessarily represent the official views of the National Institutes of Health: University of Florida Clinical and Translational Science Institute (UL1TR001427). Vanderbilt University/Vanderbilt University Medical Center: Mouse Metabolic Phenotyping Core, Vanderbilt University (DK135073 and DK020593); Hematopoietic Malignancies Biospecimen Repository; Flow Cytometry Shared Resource supported by VICC (P30 CA68485); the Vanderbilt Digestive Disease Research Center (DK058404); Cell Imaging Shared Resource (CA68485, DK20593, DK59637, EY08126); and the High-throughput Screening (HTS) core facility supported the Vanderbilt Institute for Chemical Biology and the Vanderbilt Ingram Cancer Center (P30 CA68485).

## Extended Data Figures

**Extended Data Figure 1 related to Figure 1.**
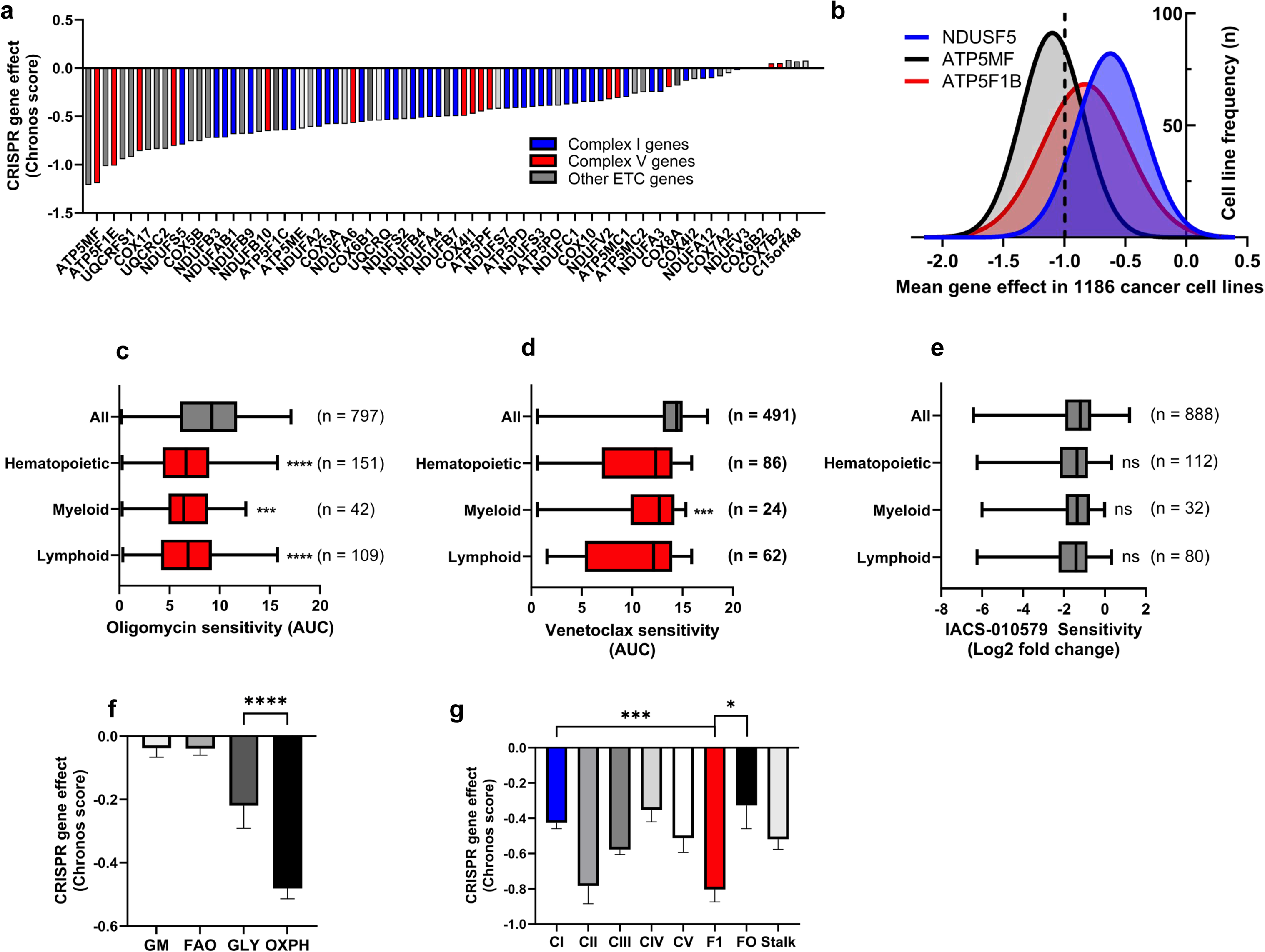
ETC complex and ATP synthase subunit gene dependencies across cancer cell lines. **(a)** CRISPR-Cas9 gene dependency scores (Chronos) for individual nuclear-encoded ETC genes ranked by mean effect across 1,186 cancer cell lines (DepMap Public 25Q2). **(b)** Distribution of mean CRISPR-Cas9 gene dependency scores (Chronos) across 1,186 cancer cell lines for three representative ETC genes: NDUSF5 (Complex I, blue), ATP5MF (Complex V, black), and ATP5F1B (Complex V, red). Dashed line indicates the pan-essentiality threshold (Chronos score = −1.0). **(c–e)** Sensitivity to oligomycin (c; area under the dose-response curve, AUC), venetoclax (d; AUC), and IACS-010759 (e; log₂ fold change) across all cancer cell lines and hematopoietic sub-lineages available in the CTD^2^ and PRISM Repurposing Public 24Q2 datasets. Comparisons between all cancer cell lines and each hematopoietic sub-lineage performed by Mann-Whitney U test. **(f)** Mean CRISPR-Cas9 gene dependency scores (Chronos ± SEM) across four metabolic gene sets — glutamine metabolism (GM), fatty acid oxidation (FAO), glycolysis (GLY), and OXPHOS — in 29 AML cell lines from DepMap Public 25Q2. Comparison between GLY and OXPHOS performed by Mann-Whitney U test. **(g)** Mean CRISPR-Cas9 gene dependency scores (Chronos ± SEM) grouped by ETC complex (CI–CV) and ATP synthase subunit class (F1, FO, peripheral stalk) in 29 AML cell lines from DepMap Public 25Q2. Comparisons performed by Mann-Whitney U test; CI vs CV and F1 vs FO comparisons indicated. NS = not significant, *p<0.05, **p<0.01, ***p<0.001.

**Extended Data Figure 2 related to Figure 1.**
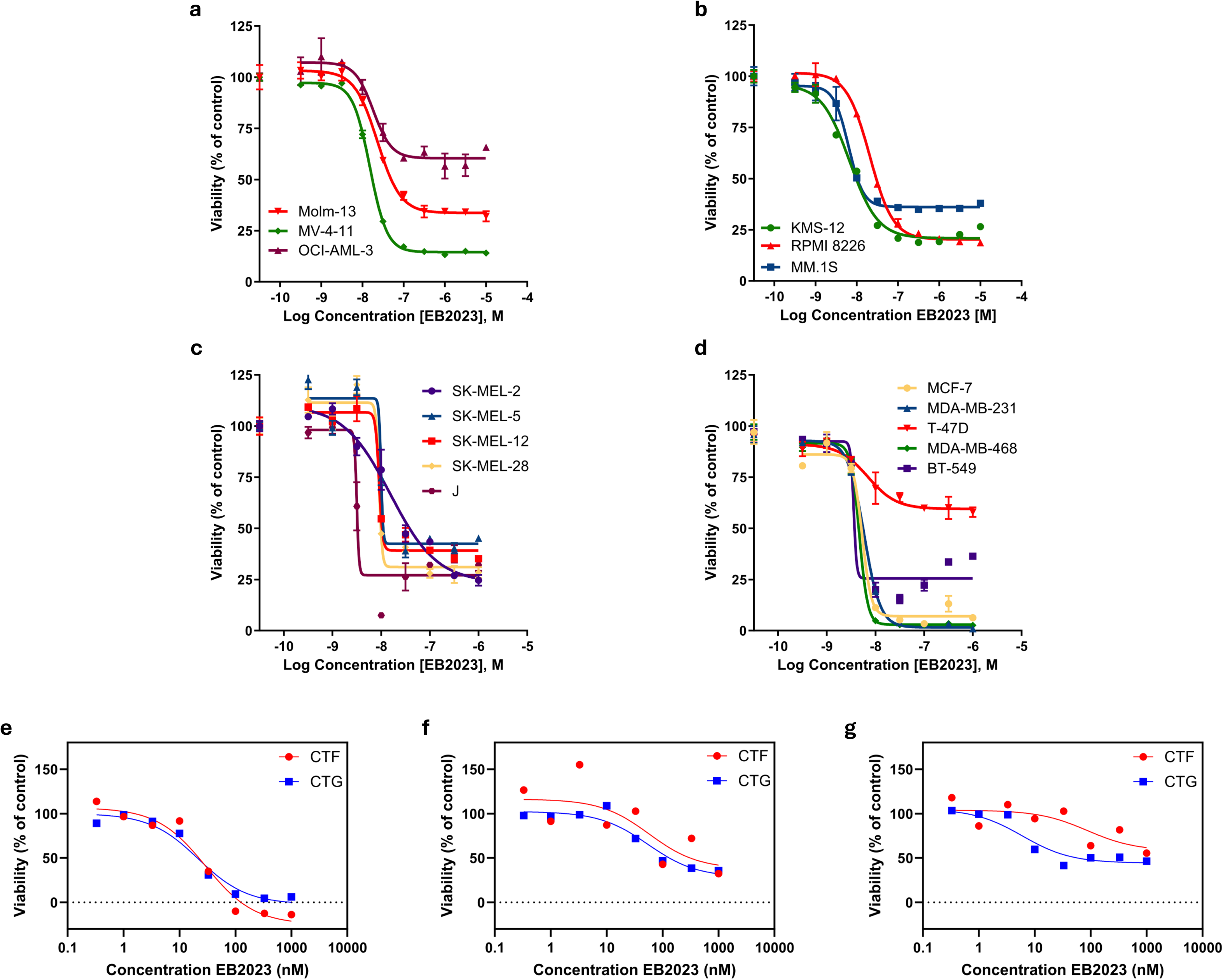
EB2023 has broad anti-cancer activity across multiple tumor lineages. (a–d) Dose-response curves for EB2023 across AML (a), multiple myeloma (b), melanoma (c), and breast cancer (d) cell lines, assessed by CellTiter-Glo viability assay at 72 hours (n = 3 replicates per condition). Curves were fit by nonlinear regression using a four-parameter logistic model. **(e–g)** Comparison of EB2023 dose-response curves measured by CellTiter-Glo (CTG, blue) and CellTiter-Fluor (CTF, red) in MV-4-11 (e), Molm-13 (f), and NB-4 (g) cells at 72 hours (n = 3 replicates per condition). Curves were fit by nonlinear regression using a four-parameter logistic model. Data shown as mean ± SEM.

**Extended Data Figure 3 related to Figure 1.**
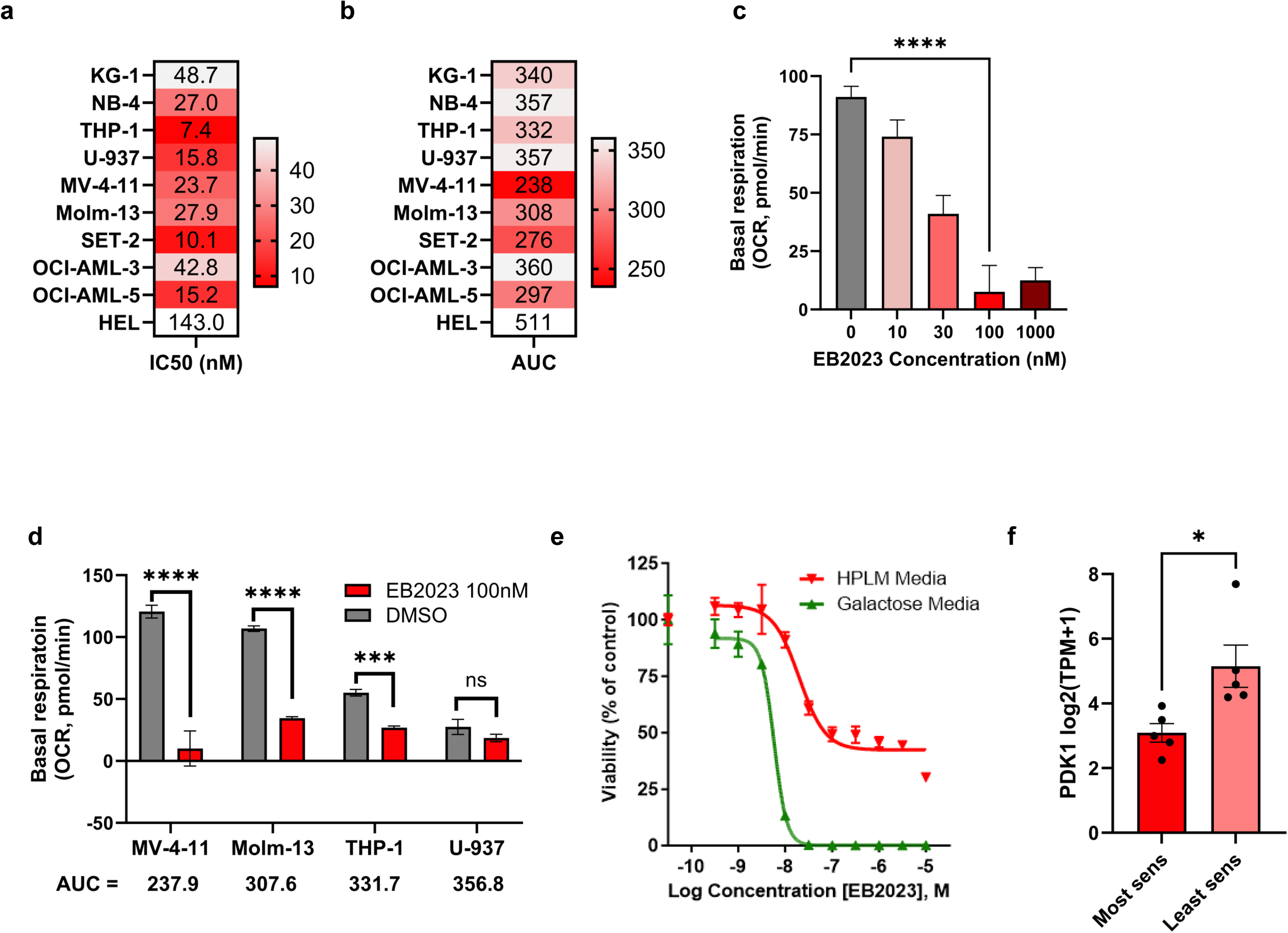
EB2023 suppresses AML respiration to affect its anti-AML activity. (a-b) IC_50_ (a, nM) and AUC (b) for 10 AML cell lines tested across an EB2023 dose-response range relative to DMSO control, assessed by CellTiter-Glo viability assay at 48 hours. **(c)** Basal respiration (OCR, pmol/min) of MV-4-11 for 24 hours, assessed by Seahorse XF Mito Stress Test (n = 3). Comparisons to vehicle performed by Welch’s unpaired t-test. **(d)** Basal respiration (OCR, pmol/min) of AML cell lines treated with EB2023 or DMSO for 24 hours, assessed by Seahorse XF Mito Stress Test (n = 3). Comparisons performed by Welch’s unpaired t-test. **(e)** Viability (CTG) of MV-4-11 cells across an EB2023 dose-response range cultured in human plasma-like medium (HPLM) or galactose-substituted medium (n = 3). Curves were fit by nonlinear regression using a four-parameter logistic model. **(f)** PDK1 expression (log₂ TPM+1) in the five most sensitive (MV-4-11, THP-1, Molm-13, OCI-AML-5, SET-2) and five least sensitive (NB-4, KG-1, U-937, OCI-AML-3, HEL) AML cell lines stratified by EB2023 AUC from CTG dose-response assay (n = 5 per group). Expression data obtained from DepMap Public 25Q2. Comparison performed by Welch’s unpaired t-test. * ns = not significant, *p<0.05, ***p<0.001, ****p<0.0001.

**Extended Data Figure 4 related to Figure 2.**
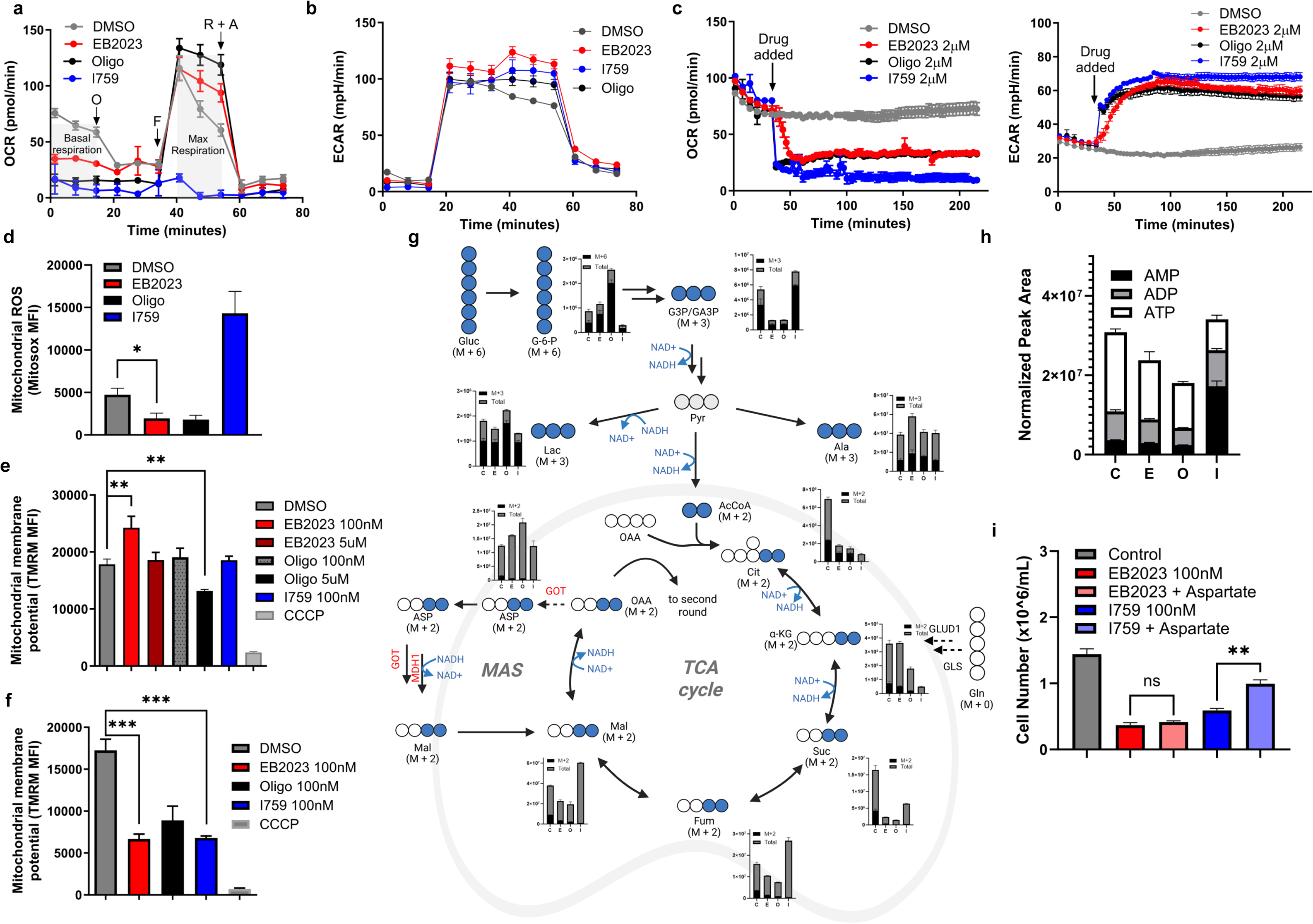
Metabolic effects of OXPHOS inhibitors in Molm-13 cells and additional mechanistic characterization in MV-4-11 cells. (a–b) OCR (a) and ECAR (b) of Molm-13 cells pretreated for 24 hours with DMSO, EB2023 (100 nM), oligomycin (100 nM), or I759 (100 nM), measured by Seahorse XF Mito Stress Test (n = 3). O, oligomycin; F, FCCP; R+A, rotenone and antimycin A. **(c)** Real-time OCR (left) and ECAR (right) of MV-4-11 cells measured by Seahorse XF analysis immediately following acute injection of drug (n = 3).**(d)** Mitochondrial ROS levels (MitoSOX MFI) in viable (Annexin V−) MV-4-11 cells treated with DMSO, EB2023 (100 nM), oligomycin (100 nM), or I759 (100 nM) for 18 hours, measured by flow cytometry (n = 3). Data shown as mean ± SEM. **(e–f)** Mitochondrial membrane potential (TMRM MFI, non-quenching concentration of 20 nM) in MV-4-11 cells treated with DMSO, EB2023 (100 nM), oligomycin (100 nM), I759 (100 nM), or CCCP (positive control) for one hour (e) or 24 hour (f), measured by flow cytometry (n = 3). **(g)** Stable isotope tracing schematic of [U-¹³C₆]-glucose metabolism in Molm-13 cells treated with DMSO (C), EB2023 (E), oligomycin (O), or I759 (I) at 100 nM for 24 hours (n = 4). Bar graphs depict labeled (M+x, black) and total metabolite abundances (grey) measured by LC-MS for glycolytic and TCA cycle intermediates. NAD⁺/NADH cofactor utilization is annotated at relevant steps. **(h)** Total abundance of ATP, ADP, and AMP in Molm-13 cells treated with DMSO (C), EB2023 (E), oligomycin (O), or I759 (I) at 100 nM for 24 hours, measured by LC-MS (n = 4). **(i)** Cell counts of MV-4-11 cells treated with DMSO, EB2023 (100 nM), EB2023 + aspartic acid (10 µM), I759 (100 nM), or I759 + aspartic acid (10 µM) for 48 hours, assessed by automated cell counting (n = 3). Data shown as mean ± SEM. Error bars represent mean ± SEM throughout. All comparisons to DMSO performed by Welch’s unpaired t-test. ns = not significant, *p<0.05, **p<0.01, ***p<0.001, ****p<0.0001.

**Extended Data Figure 5 related to Figure 3.**
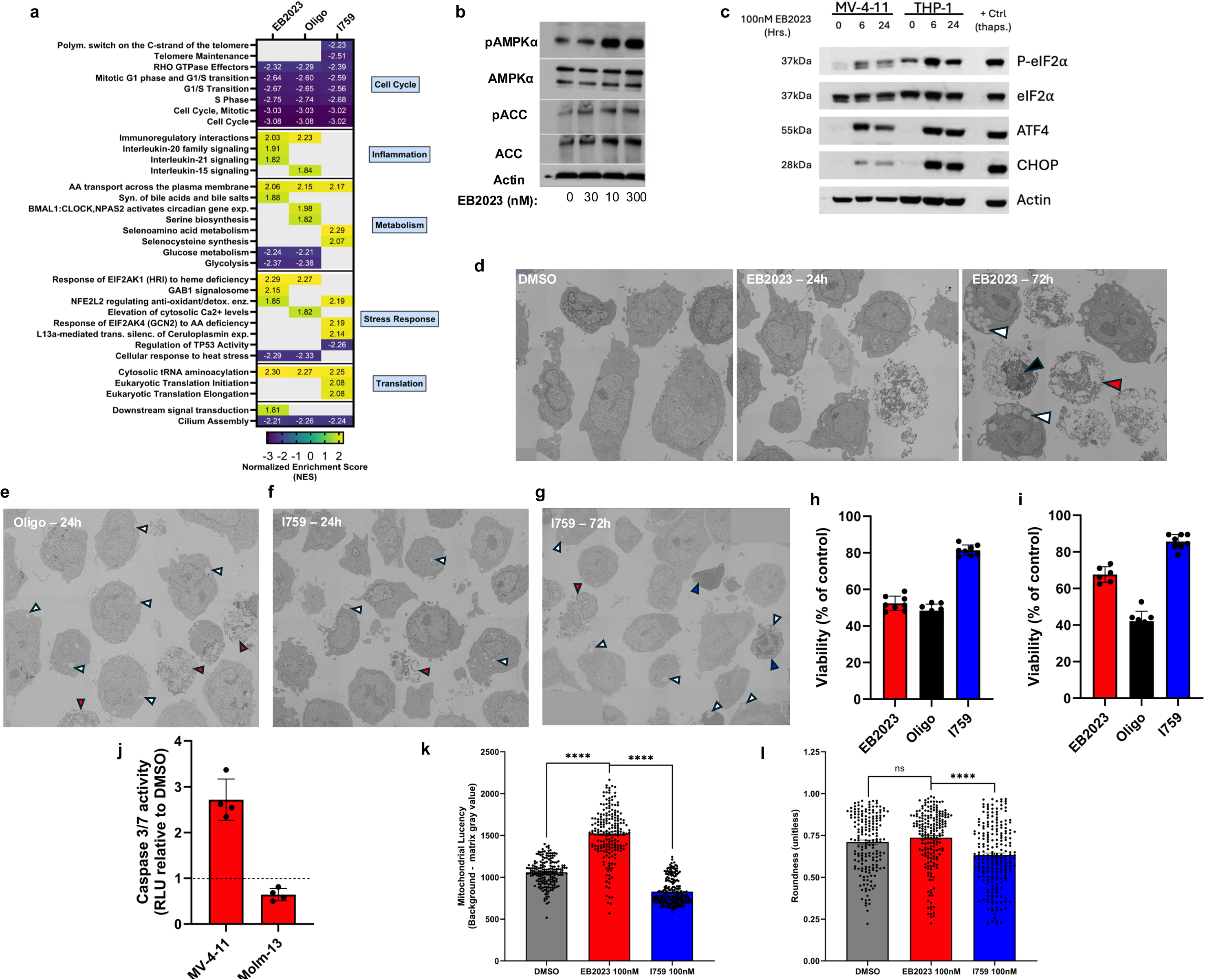
Stress responses to OXPHOS inhibition in MV-4-11 and THP-1 cells. **(a)** Heatmap of normalized enrichment scores (NES) from GSEA of the top Reactome gene sets per comparison in THP-1 cells treated with EB2023, oligomycin, or I759 at 100 nM for 24 hours versus DMSO. Gene sets are grouped by biological category. Positive NES (warm colors) indicates upregulation; negative NES (cool colors) indicates downregulation relative to DMSO. **(b)** Immunoblot analysis of phospho-AMPKα (Thr172), total AMPKα, phospho-ACC (Ser79), total ACC, and actin in MV-4-11 cells treated with EB2023 for 24 hours. **(c)** Immunoblot analysis of phospho-eIF2α (Ser51), total eIF2α, ATF4, CHOP, and actin in MV-4-11 and THP-1 cells treated with EB2023 (100 nM) for 0, 6, or 24 hours. Thapsigargin-treated MV-4-11 cells serve as a positive control for ISR activation. **(d)** Representative TEM images of MV-4-11 cells treated with DMSO or EB2023 (100 nM) for 24 or 72 hours. White arrowheads indicate cytoplasmic vacuolization; red arrowheads indicate loss of plasma membrane integrity consistent with necrotic morphology; black arrowheads indicate nuclear changes. **(e–g)** TEM images of MV-4-11 cells treated with oligomycin (100 nM) for 24 hours (e), I759 (100 nM) for 24 hours (f), or I759 (100 nM) for 72 hours (g). White arrowheads indicate cytoplasmic vacuolization; red arrowheads indicate loss of plasma membrane integrity consistent with necrotic morphology; blue arrowheads indicate condensed cells with apoptotic nuclear features including condensed and marginalized chromatin. **(h–i)** Relative viability (% trypan blue-negative cells normalized to DMSO) of MV-4-11 (h) and Molm-13 (i) cells treated with EB2023 (100 nM), oligomycin (100 nM), or I759 (100 nM) for 24 hours (n = 3). **(j)** Caspase 3/7 activity (relative light units normalized to DMSO; dashed line indicates DMSO reference) in MV-4-11 and Molm-13 cells treated with EB2023 (100 nM) for 24 hours, measured by Caspase-Glo assay (n = 3). **(k)** Mitochondrial lucency (background − matrix gray value) quantified by ImageJ analysis of TEM images from MV-4-11 cells treated with DMSO, EB2023 (100 nM), or I759 (100 nM) for 24 hours. Higher values indicate greater electron lucency of the mitochondrial matrix. n = 200 mitochondria per condition from an unbiased representative selection of TEM images. Comparisons performed by Welch’s unpaired t-test. **(l)** Mitochondrial roundness (unitless, 0–1 scale; 1 = perfect circle) quantified by ImageJ analysis of the same TEM image dataset as panel k. Comparisons performed by Welch’s unpaired t-test. All bar graphs represent mean ± SEM. ns = not significant, ****p<0.0001.

**Extended Data Figure 6 related to Figure 5.**
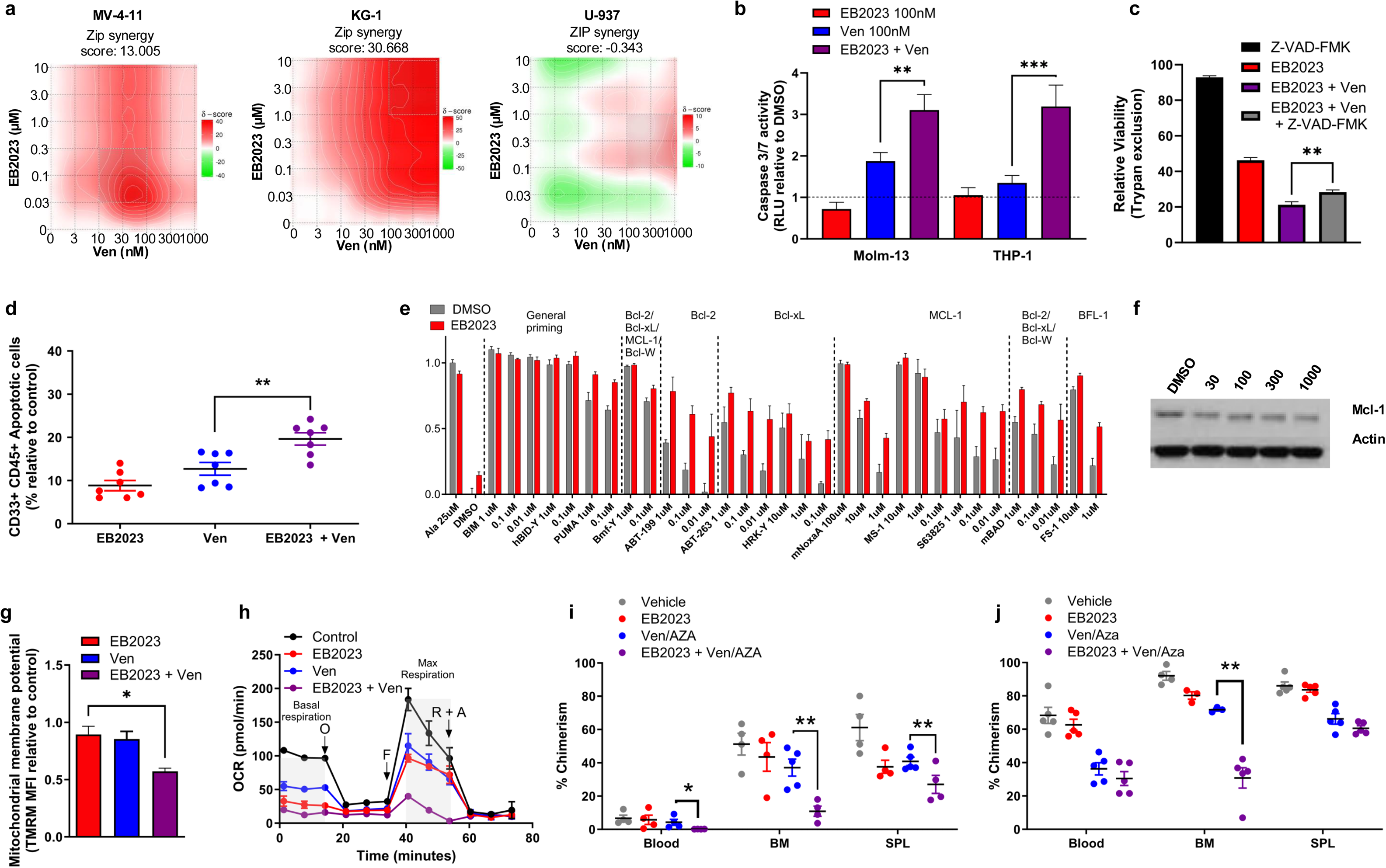
EB2023 and venetoclax combination induces caspase-dependent apoptosis and overcomes venetoclax resistance in vitro and in vivo. **(a)** ZIP synergy score matrices for EB2023 (10 nM–10 µM) and venetoclax (1 nM–1 µM) in MV-4-11, KG-1, and U-937 AML cell lines, assessed by CTG at 48 hours. ZIP synergy scores are indicated; warm colors indicate synergy, cool colors indicate antagonism (ZIP score >10 indicates synergy).^37^ **(b)** Caspase 3/7 activity (relative light units normalized to DMSO; dashed line indicates DMSO reference) in Molm-13 and THP-1 cells treated with EB2023, venetoclax, or their combination for 24 hours (n = 3). **(c)** Relative viability (trypan blue exclusion) of MV-4-11 cells treated for 48 hours (n = 3). **(d)** Percentage of apoptotic AML blasts (CD33+/CD45+ Annexin V+/PI−) in primary AML patient leukapheresis samples treated 18 hours (n = 8). **(e)** Extended BH3 profiling panel from THP-1 cells pretreated with DMSO or EB2023 (100 nM) for 24 hours (n = 3). **(f)** Immunoblot of MCL1 and actin in MOLM-13 cells treated for 24 hours. **(g)** Relative mitochondrial membrane potential (TMRM MFI normalized to DMSO) in viable (Annexin V−) THP-1 cells treated with indicated conditions for 30 minutes (n = 3). **(h)** OCR of THP-1 cells treated with indicated conditions for 24 hours by Seahorse XF Mito Stress Test (n = 3). O, oligomycin; F, FCCP; R+A, rotenone and antimycin A. **(i–j)** Human AML chimerism (% hCD45+ cells) in blood, BM, and SPL of MV-4-11 CDX (i) and AML13 PDX (j) NSGS mice (n = 6 per group) treated with vehicle, EB2023 (0.1mg/kg), Ven/AZA (venetoclax 20 mg/kg, azacitidine 1.5 mg/kg), or EB2023 + Ven/AZA. All bar and dot plots represent mean ± SEM. Comparisons performed by Welch’s unpaired t-test. ns = not significant, *p<0.05, **p<0.01, ***p<0.001.

**Extended Data Figure 7 Related to Figure 5.**
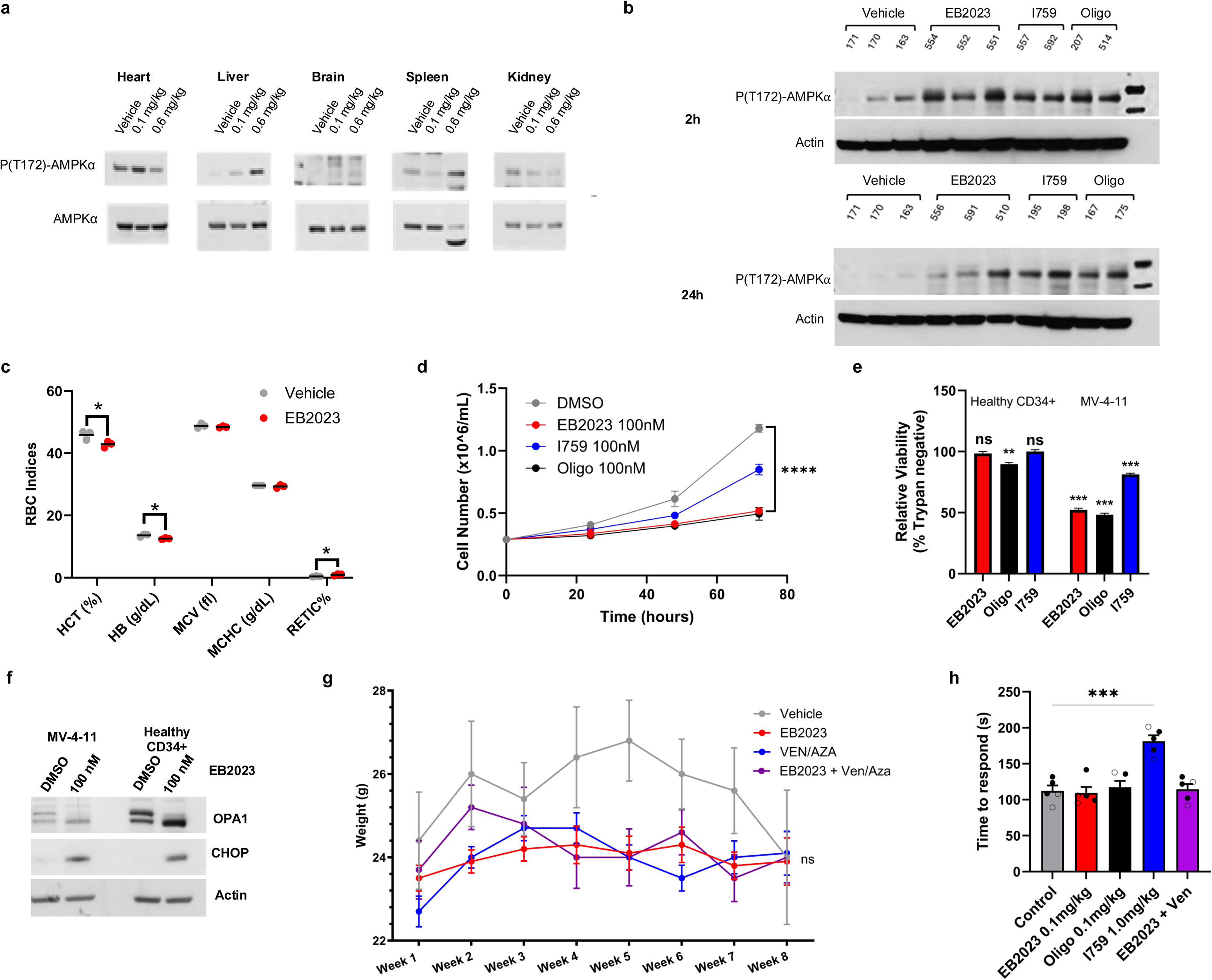
EB2023 effects on normal tissue *in vitro* and *in vivo*. **(a)** Immunoblot analysis of phospho-AMPKα (Thr172) and total AMPKα in heart, liver, brain, spleen, and kidney harvested by freeze-clamp 2 hours after a single dose from NSGS mice treated with vehicle or EB2023 (0.1 or 0.6 mg/kg). **(b)** Immunoblot analysis of phospho-AMPKα (Thr172) and actin in bone marrow cells from MV-4-11 CDX NSGS mice treated with vehicle, EB2023, I759, or oligomycin and harvested at 2 and 24 hours post-dose (n = 2–3 per group per timepoint). Individual animal identifiers are indicated above each lane. **(c)** Red blood cell indices — hematocrit (HCT), hemoglobin (HB), mean corpuscular volume (MCV), mean corpuscular hemoglobin concentration (MCHC), and reticulocyte percentage (RETIC%) — measured by automated cytometry in CD1 mice treated with vehicle or EB2023 (0.1 mg/kg, once daily, 5 days on/2 days off) for 2 weeks (n = 4 per group). **(d)** Cell counts of healthy CD34+ umbilical cord blood HSPCs cultured in StemSpan II expansion medium and treated for 72 hours (n = 3). Statistical comparisons to DMSO performed by two-way repeated measures ANOVA with Dunnett’s post-hoc test. **(e)** Relative viability of healthy CD34^+^ umbilical cord blood HSPCs and MV-4-11 cells treated with EB2023, oligomycin, or I759 at 100 nM for 48 hours (n = 4). **(f)** Immunoblot analysis of OPA1, CHOP, and actin in MV-4-11 cells and healthy CD34+ HSPCs treated for 24 hours. **(g)** Body weight of AML06 PDX NSGS mice during continuous treatment with vehicle, EB2023, Ven/AZA, or EB2023 + Ven/AZA (n = 6 per group). Data shown as mean ± SEM. Body weight was compared across treatment groups over 8 weeks by two-way ANOVA. The treatment × time interaction was not significant (F(21,248)=0.76, p=0.772), indicating no differential weight change over the treatment period. ns = not significant. **(h)** Time to respond (seconds) in adhesive removal test in CD1 mice treated with indicated conditions (n = 5 per group). Comparisons to control performed by one-way ANOVA with Tukey’s post-hoc test. All bar and dot plots represent mean ± SEM. Comparisons performed by Welch’s unpaired t-test unless otherwise stated. *p<0.05, ****p<0.0001.

