## Supplemental files for "Specific F_1_ ATP synthase inhibition delivers transient mitochondrial stress for selective targeting of acute myeloid leukemia"

### Supplementary Table 1: AML Patient Sample Clinical and Molecular Information

Supplementary Table 7: AML Patient Sample Clinical Information

| Label | Sample ID | FAB Subtype | Age | WBC | Blasts (%) | NGS | Karyotype | Cytogenetics | Source |
|---|---|---|---|---|---|---|---|---|---|
| AML 01 | 22-01-016 | M5 | 49 | 174.8 | 65 | FLT3-ITD | 46,XY[20] | Neg FISH | Leukapheresis |
| AML 02 | 21-06-001 | NA | 69 | 408.5 | 57 | FLT3-ITD, DNMT3A, TP53 | Unknown | NA | Leukapheresis |
| AML 03 | 19-06-004 | M5 | 77 | 44.2 | 66 | TET2x2, TP53 | 44,XX,der(4)t(4;17)(q31;q21),-5,+8,-9,-17[19]/46,XX[1] | Trisomy 8, del17p | Leukapheresis |
| AML 04 | 22-05-002 | M5 | 42 | 235.6 | 49 | KRAS x2, GATA2 x2 | 46,XY,t(11;19)(q23;p13.1)[17]/46,XY[3] | KMT2Ar t(11;19) | Leukapheresis |
| AML 05 | 22-01-020 | M5 | 75 | 249 | 89 | FLT3-ITD, RUNX1, U2AF1 | 46,XY[20] | Neg FISH | Leukapheresis |
| AML 06 | 21-12-019 | NA | 63 | 137.1 | 67 | NPM1, FLT3-ITD, DNMT3A, ASXL1 | 46,XX[20] | Neg FISH | Leukapheresis |
| AML 07 | 15-08-003 | M5 | 53 | 4.5 | 50 | NPM1, FLT3-TKD, DNMT3A | 47~48,XY,t(3;4)(q25;q21),+9,del(20(q11.2q13.3),+21[cp16]/50,idem,+Y,add(1)(q32),del(6)(q13),add(7) (q32),+10,+13,-16,+22[4] | NA | Bone Marrow |
| AML 08 | 24-01-002F | M4/M5 | 77 | 58.3 | 90 | FLT3, NPM1, DNMT3A, PTPN11 x3, TET2 x2 | 46,XY[20] | Neg FISH | Bone Marrow |
| AML 09 | 23-02-010F | NA | 80 | 65.5 | 85 | RUNX1, TET2x2, IDH2, JAK2, SRSF2 | 47,XY,+8[4]/46,XY,add(21)(q22)[4]/46,XY[12] | Trisomy 8 | Bone Marrow |
| 0 | 20-08-009F | NA | 79 | 48.9 | 89 | ASXL1, RUNX1, PTPN11 x2 | 48,XY,+8,+21[19]/46,XY[1] | Trisomy 8 | Bone Marrow |
| AML 11 | 24-02-005F | NA | 71 | 81.2 | 77 | FLT3 x2, NPM1, DNMT3A, WT1 | 46,XY[20] | Neg FISH | Bone Marrow |
| AML 12 | 23-09-011F | M5 | 62 | 55 | 88 | FLT3, NPM1, DNMT3A, TET2, ZRSR2 | 46,XY[20] | Neg FISH | Bone Marrow |
| AML 13 | 17-12-011 | M5 | 65 | 165.2 | 95 | FLT3-TKD, TET2x2, TP53x2 | 45,XX,-16,add(17)(p11.2)[18]/46,XX[2] | Loss of 16q22 | Leukapheresis |
| AML 14 | 18-12-001 | NA | 62 | 62.9 | 94 | NPM1, FLT3-ITD, IDH2, DNMT3A, PTPN11 | 46,XX[20] | Neg FISH | Leukapheresis |
| AML 15 | 16-05-017 | NA | 51 | 65.5 | 86 | FLT3-ITD, NPM1, DNMT3A, WT1 | 46,XX,?der(17)del(17)(p13)inv(17)(p13q11.2)[11]/46,idem,del(12)(p11.2p13)[9] | NA | Leukapheresis |

**Supplemental Table 1. AML patient sample clinical and molecular information.**

Annotated AML samples with clinical, laboratory and molecular data at time of sample acquisition.

#### Supplementary Table 2: EB2023 PK studies in CD1 mice

| Parameter | IV | IP | PO |
|---|---|---|---|
| Dose (mg/kg) | 0.17 | 0.17 | 0.17 |
| $T_{1/2}$ (h) | 0.5 | - | - |
| $C_0$ / $C_{max}$ (ng/mL) | 71.5 | 50.9 | - |
| $AUC_{all}$ (h*ng/mL) | 50.6 | 42.9 | - |
| $AUC_{INF\_obs}$ (h*ng/mL) | 53.3 | 44.6 | - |
| Bioavailability (%) | - | 83.7 | 0.0 |

**Supplemental Table 2. EB2023 and pharmacodynamic studies.**

Pharmacokinetic parameters calculated from serially measured EB2023 plasma concentrations as determined by LC-MS/MS in CD-1 mice treated with a single dose of EB2023 (0.170 mg/kg) by either IP, IV or PO routes. $C_0$ = extrapolated initial plasma peak concentration (IV), $C_{max}$ = plasma peak concentration, $T_{1/2}$ = elimination half-life, AUC = area under the concentration-time curve.

#### Supplemental Figure 1: OXPHOS inhibitor dose response comparisons in primary AML patient samples

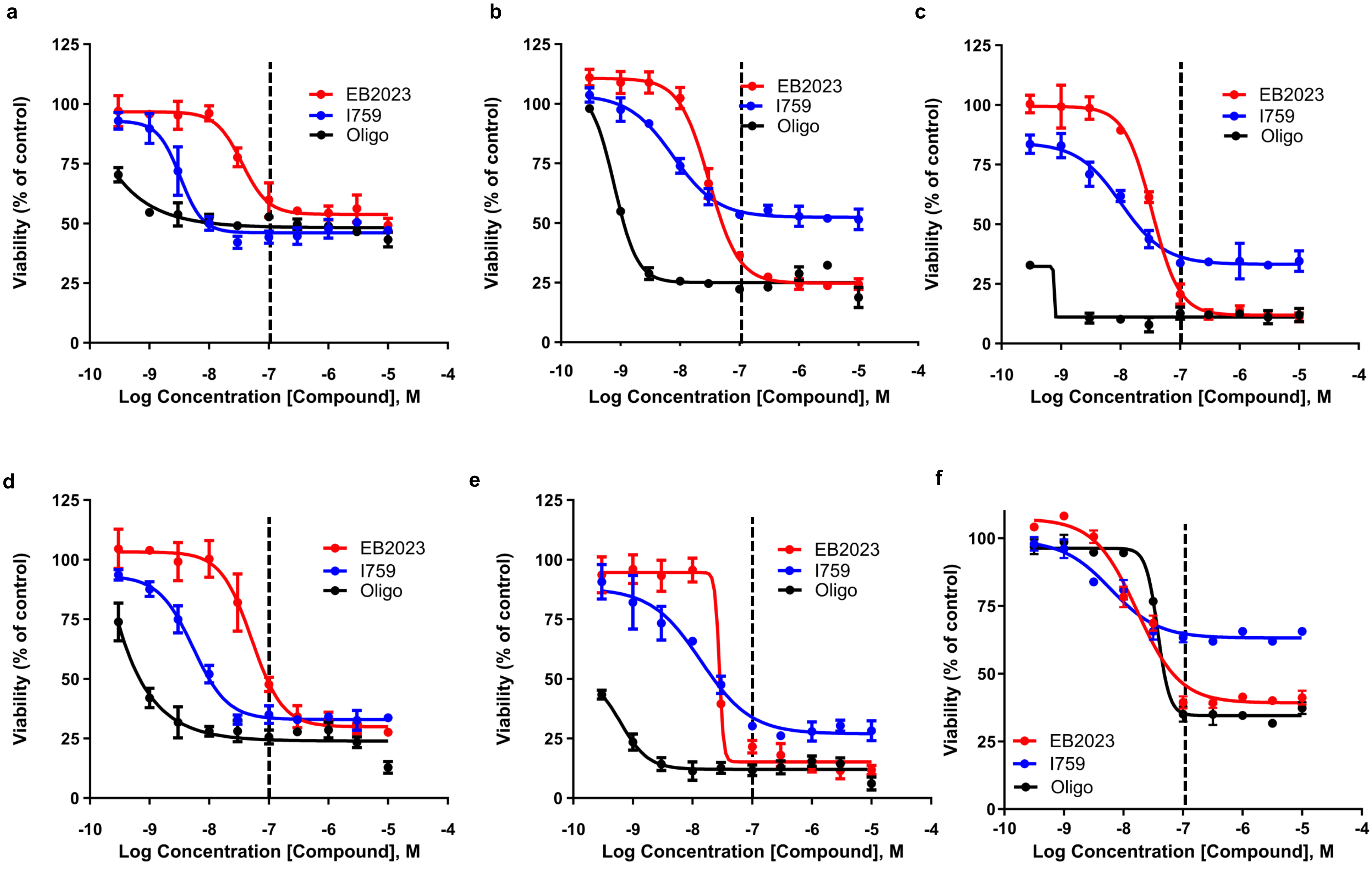

**Supplemental Figure 1. OXPHOS inhibitor dose-response comparisons in primary AML patient samples.** **(a–e)** Dose-response curves for EB2023, IACS-010759 (I759), and oligomycin in primary AML patient samples AML08 (a), AML09 (b), AML10 (c), AML11 (d), and AML12 (e), assessed by CellTiter-Glo viability assay at 72 hours (n = 3 replicates per condition). Dashed vertical line indicates 100 nM reference concentration. **(f)** Dose-response curves for EB2023, I759, and oligomycin in Molm-13 cells assessed by CellTiter-Glo at 72 hours (n = 3 replicates per condition). Curves were fit by nonlinear regression using a four-parameter logistic model. Data shown as mean ± SEM.

#### Supplemental Figure 2. Calorimetry assessment of ATP synthase inhibitors in CD1 mice.

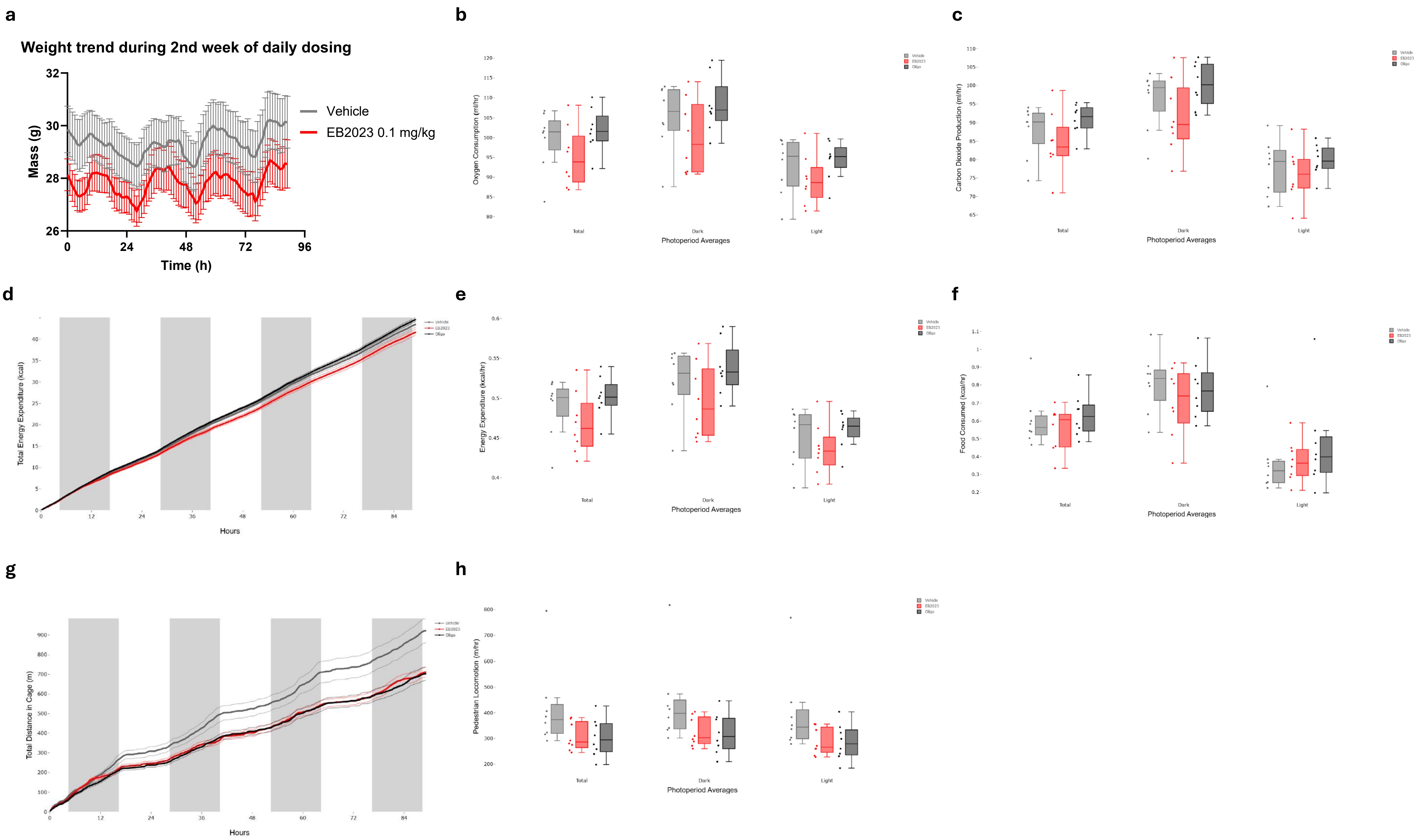

**Supplemental Figure 2. Indirect calorimetry analysis of CD1 mice treated with vehicle or EB2023 during the second week of daily dosing. (a)** Body mass trend over 96 hours of measurement. **(b–c)** Oxygen consumption ($VO_2$, ml/hr) and carbon dioxide production ($VCO_2$, ml/hr) averaged over total, dark, and light photoperiods. **(d)** Cumulative total energy expenditure (kcal) over 96 hours. **(e)** Energy expenditure (kcal/hr) averaged over total, dark, and light photoperiods. **(f)** Food consumed (kcal/hr) averaged over total, dark, and light photoperiods. **(g)** Cumulative total distance traveled in cage (m) over 96 hours. **(h)** Ambulatory locomotor activity (pedometer, m/hr) averaged over total, dark, and light photoperiods. Data shown as median with interquartile range (n = 8 per group). Mass-dependent variables ($VO_2$, $VCO_2$, energy expenditure, food intake) were analyzed by ANCOVA with body mass as covariate; mass-independent variables (locomotor activity) were analyzed by ANOVA. Statistical analysis performed using CalR2 (Mina et al., *Cell Metabolism* 2018).
